# SOX2-induced IL1α-mediated immune suppression drives epithelial dysplasia malignant transformation

**DOI:** 10.1101/2024.12.06.626475

**Authors:** Hülya F. Taner, Wang Gong, Zackary R. Fitzsimonds, Zaiye Li, Yuesong Wu, Yumin He, Kohei Okuyama, Wanqing Cheng, Jung Kuczura, Sashider Rajesh, Andriana Manousidaki, Shuo Feng, Miki Lee, Felipe Nör, Emily Lanzel, Shadmehr Demehri, Peter J. Polverini, Jacques E. Nör, Thomas D. Wang, Jianwen Que, Haitao Wen, Yuying Xie, James J. Moon, Yu Leo Lei

## Abstract

Squamous cell carcinomas (SCC) are often preceded by potentially malignant precursor lesions, most of which remain benign. The terminal exhaustion phenotypes of effector T-cells and the accumulation of myeloid-derived suppressor cells (MDSC) have been thoroughly characterized in established SCC. However, it is unclear what precancerous lesions harbor a bona fide high risk for malignant transformation and how precancerous epithelial dysplasia drives the immune system to the point of no return. Here we show that expression of SRY-box transcription factor 2 (SOX2) in precancerous lesions imparts an irreversible risk that recruits suppressive myeloid cells by promoting the release of CCL2. We developed a unique genetically engineered mouse model (GEMM) to recapitulate the malignant transformation of epithelial dysplasia to SCC in the oral mucosa with high histologic and phenotypic fidelity. Using a combination of longitudinal human specimens and the Sox2-GEMM, we found that the myeloid cells in precancerous epithelial dysplasia exhibit a distinctive dichotomous profile featuring high levels of IL-1α-SLC2A1 and low levels of type-I interferon (IFN-I) signatures, which occurs before SCC emerges histologically. Brief priming of myeloid cells with IL-1α desensitizes them to IFN-I agonists and makes myeloid-derived suppressor cells (MDSC) even more suppressive of T-cell activation. Mechanistically, IL-1 activation represses the expression of DHHC3/7 enzymes, which are responsible for the palmitoylation of stimulator of interferon genes (STING). Early blockade of IL1 signaling using pharmacologic and genetic approaches similarly reduces MDSC and SLC2A1^high^ myeloid cells, suppresses epithelial dysplasia transformation, and extends survival. This work establishes a previously unrecognized SOX2-CCL2-IL1 pathway that leads to irreversible immune escape when precancerous epithelial lesions transform.

## Main

Extensive characterization of the microenvironment of established cancers has revealed complex compensatory immune suppressive mechanisms, such as the engagement of multiple immune checkpoint receptor pathways, massive recruitment of myeloid-derived suppressor cells (MDSC), and the expansion of regulatory T-cells and suppressive B-cell subsets. Thus, targeting a single pathway of immune suppression at late stages is often insufficient. Human papillomavirus (HPV)^-^ Head and Neck Squamous Cell Carcinoma (HNSCC) is such an example, which shows a modest response rate to immune checkpoint inhibitors (ICI) despite high tumor mutational burden ^1, 2^. HPV^-^ HNSCC is preceded by pre-cancerous lesions, coined as oral epithelial dysplasia (OED). However, it remains unclear how intra-lesional immune infiltrate reaches a point of no return to terminal exhaustion during malignant transformation. OED is among the most common mucosal lesions, affecting ∼6% of the global population, with an estimated 12% transformation rate. Despite vigilant monitoring, a subset of OED transforms into HNSCC. Surgical excision cannot reverse field cancerization. The identification of the earliest events leading to immune tolerance at the transformation site offers promise to develop better immunoprevention strategies for high-risk lesions.

Whole exome sequencing of squamous cell carcinomas of the lung, esophagus, skin, and head and neck has revealed that the amplification of the 3q26.3 locus is a common hotspot driver event. Two genes at this locus, *SOX2* and *PIK3CA*, are amplified in about 20% of HNSCC^3^. Patients with Fanconi anemia suffer from an extremely early onset of HNSCC with some cases seen in patients in the first to second decades of their lives, which provides an opportunity to identify high risk events for epithelial malignant transformation. The HNSCC cases associated with Fanconi anemia have much fewer point mutations but harbor *SOX2* amplification in 78% of the cases^4^, further highlighting the important role of this driver event. Previously, we found that SOX2-expressing tumors excluded cytotoxic T-lymphocytes (CTL) from the tumor microenvironment (TME) in an implantable syngeneic mouse model^5^. However, how *SOX2* amplification in transforming epithelial cells establishes peripheral immune tolerance is largely unknown.

Here we developed a genetically engineered mouse model (GEMM) mimicking *SOX2* amplification in the oral epithelium and utilized matched longitudinal human specimens to identify early high-risk features, which potentiate the irreversible malignant transformation of OED. We have revealed unique early immunometabolic markers arising in the pre-cancerous lesions prior to the appearance of HNSCC histology. SOX2 promotes CCL2 to expand intra-lesional myeloid cells exhibiting high expression levels of IL-1 signatures, glycolysis signatures, and low expression levels of type-I interferon (IFN-I) signatures. As a mechanism, IL-1α priming desensitizes myeloid cells to cytoplasmic DNA-induced IFN-I activation by inhibiting the palmitoylation of the adaptor protein, Stimulator of interferon genes (STING). Early blockade of IL1 signaling through genetic or pharmacologic approaches delays the onset of SOX2-driven epithelial malignant transformation.

### Oral epithelial cell expression of Sox2 drives HNSCC initiation

To longitudinally monitor precancer transformation, we crossed *K5-CreER* with *Rosa26-Sox2-Enhanced Green Fluorescence Protein (EGFP)* and topically painted the oral mucosa with tamoxifen to produce the Sox2-GEMM, in which oral epithelial cells expressed Sox2-EGFP (Figure 1A). We generated heterozygous and homozygous compound strains and noted a dose-dependent expression of *Sox2* in the oral epithelium (Figure 1B). Human HNSCC contains diagnostic features, including high-grade cytologic features, the transition between normal and dysplastic epithelium, the presence of invasive tumor islands, the formation of keratin pearls, and the establishment of stroma and immune-cell infiltration (Figure 1C and 1D). However, the implantation-based models cannot recapitulate all key histologic features. For example, human patient-derived xenograft (PDX) retains many high-grade cytologic features and abundant keratin. However, these implantable tumors often grow like an epidermoid cyst in vivo with keratin being the most abundant content of the cyst-like structures (Figure 1E and 1F). More importantly, PDX cannot model tumor-immune interactions. Multiple syngeneic HNSCC models have been established, which provide valuable tools to study the TME. However, due to the nature of these models, they cannot recapitulate the early epithelial transformation phase of the disease. Histologic examinations revealed spindle-cell or squamous-like high-grade tumor cells, with little stromal complexity and architectural resemblance to human HNSCC (Figure 1G and 1H).

**Figure 1.**
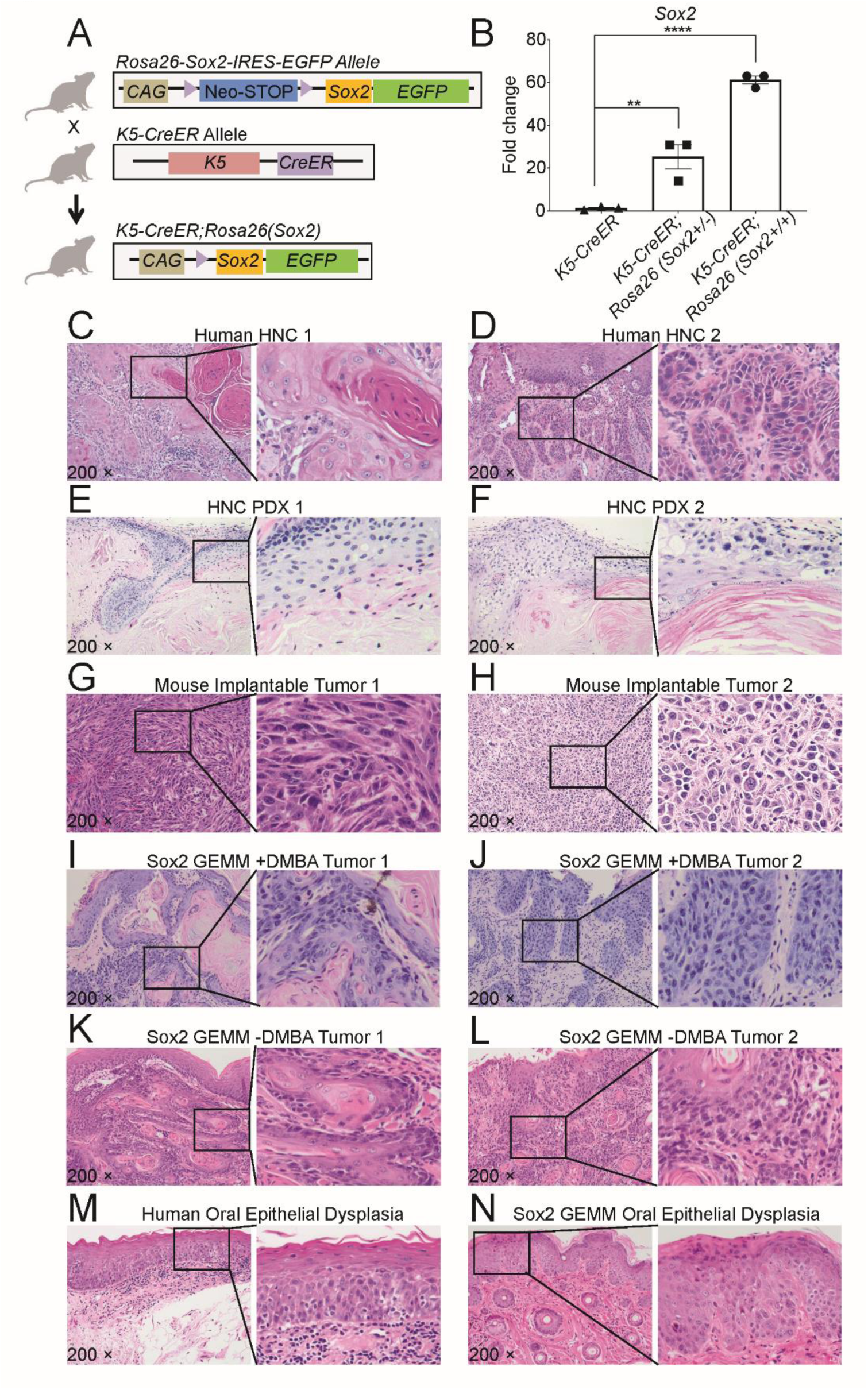
Sox2-GEMM recapitulates the stromal complexity and architectural features of human SOX2+ oral cancers. (A) Schematic depicting the generation of *KRT5-CreER;Rosa26(Sox2)* mice. (B) qPCR analysis for *Sox2* expression in buccal mucosa tissue harvested on day 7 post topical tamoxifen administration (n=3). (C-D) H&E staining of human Head and Neck Cancer (HNC) lesions. (E-F) H&E staining of human patient-derived xenograft (PDX) lesions. (G) H&E staining of HPV16 E6/E7 mEER subcutaneously implanted tumors in C57BL6/J mouse. (H) H&E staining of MOC2-EV subcutaneously implanted tumor in a C57BL6/J mouse. (I) H&E staining of extensive cancer in the buccal mucosa of Sox2-GEMM (*K5-CreER;Sox2^+/+^*^)^ mouse treated with the carcinogen 7,12-Dimethylbenz[a]anthracene (DMBA) and tamoxifen (250 mg/kg) via intraperitoneal injection (i.p.), at day 36 post administration of tamoxifen. (J) H&E staining of cancer in the buccal mucosa of a Sox2-GEMM mouse treated with DMBA and tamoxifen (75mg/kg) (i.p.), at day 26 post administration of tamoxifen. (K-L) H&E staining of cancer in the buccal mucosa of Sox2-GEMM mice treated topically with only tamoxifen (40 mg/mL). (M) H&E staining of human oral epithelial dysplasia lesion. (N) H&E staining of oral epithelial dysplasia in the buccal mucosa of a Sox2-GEMM mouse treated orally with topical administration of tamoxifen (40 mg/mL).

We treated the Sox2-GEMM or *K5-CreER* control mice with a carcinogen, DMBA, or PBS. Buccal mucosa and tongue were subjected to histologic examination. We found that Sox2 expression was a potent driver of HNSCC. The homozygous compound strain developed HNSCC with or without additional DMBA painting. The histologic features of Sox2-GEMM show high-fidelity recapitulation of human HNSCC (Figure 1I to 1L). Importantly, longitudinal monitoring of the Sox2-GEMM revealed an early transitional phase where oral epithelium exhibited dysplastic features that were highly similar to human OED, such as architectural disturbance, cell crowding, nuclear hyperchromasia, and increased mitotic counts above the basal layer (Figure 1M and 1N).

### Sox2-GEMM shows key immunophenotypic features of initiating HNSCC

The *K5-CreER^+^;Sox2^+/-^* strain without DMBA showed about 60-80% HNSCC penetrance 160 days post-induction, and the *K5-CreER^+^;Sox2^+/+^* strain showed 100% spontaneous penetrance about four to six weeks post-induction (Figure 2A). The majority of the lesions arose in the buccal mucosa. We did not observe cancers arising in the tongue without DMBA treatment. About 60% of the *K5-CreER^+^;Sox2^+/+^* mice developed tongue cancer when co-treated with DMBA. OED was seen primarily in the buccal mucosa or tongue of the *K5-CreER^+^;Sox2^+/-^*mice (Figure 2A). While the majority of the HNSCC in *K5-CreER^+^;Sox2^+/+^* mice were microscopic due to their rapid onset, some tumors were seen grossly (Figure 2B). We engineered an endomicroscope for direct laser confocal imaging and revealed EGFP-expressing Sox2^+^ transformed oral epithelial cells in live animals (Figure 2C). Then, we harvested buccal mucosa from *K5-CreER^+^;Sox2^+/+^* mice and *K5-CreER* control mice five weeks after tamoxifen was painted in the oral mucosa. We stained the paraffin-embedded oral mucosa with immunohistochemical markers and found that the invasive squamous cell carcinoma islands showed diffuse strong nuclear staining of Sox2, as expected. These tumors showed a high Ki-67 index, with the staining patterns of p63 and p53 similar to those of human HNSCC (Figure 2D).

**Figure 2.**
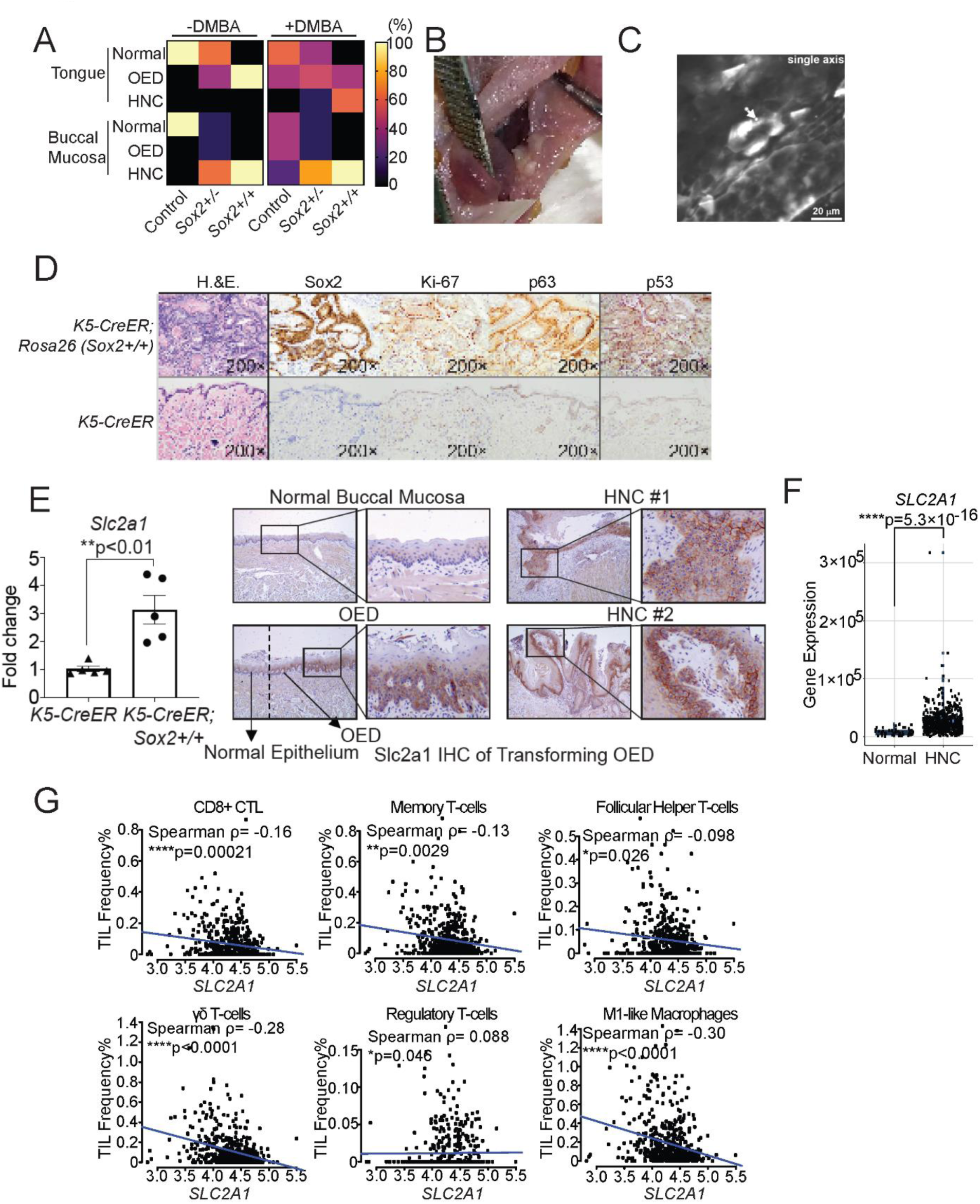
Sox2 drives squamous cell carcinomas development in the oral cavity. (A) After topical tamoxifen induction, the occurrence rates of OED and HNSCC in *K5-CreER;Sox2^+/+^*, *K5-CreER;Sox2^+/-^*or *K5-CreER* control mice treated with or without 7,12-Dimethylbenz[a]anthracene (DMBA) carcinogen are shown. Ten *K5-CreER;Sox2^+/+^*mice were treated with DMBA (n=5) or vehicle only (n=5). Sixteen *K5-CreER;Sox2^+/-^* mice were treated with DMBA (n=8) or vehicle only (n=8). Ten *K5-CreER* control mice were treated with DMBA (n=5) or vehicle only (n=5). Mice were euthanized about 4 weeks post-tamoxifen treatment. Tongue and buccal mucosa were harvested and subjected to histologic examination by two pathologists. (B) A representative Sox2-driven HNSCC on the buccal mucosa is shown. (C) Four weeks post topical tamoxifen induction, buccal mucosa from *K5-CreER;Sox2^+/+^* mice was harvested and cleared using the PEGASOS method. Representative confocal imaging examination shows Sox2^+^GFP^+^ tumor growth. (D) Buccal mucosa of tamoxifen-induced K5-CreER;Sox2^+/+^ and littermate control mice were harvested 4 weeks post-induction and subjected to IHC staining of the indicated markers.(E) Increased *Slc2a1* expression was observed in buccal mucosa of control (n=5) or Sox2^+/+^ mice (n=5) that have been treated with tamoxifen. Mice were euthanized ∼3 weeks post tamoxifen treatment and RNA was isolated from buccal mucosa for qPCR and IHC. (F) *SLC2A1* expression levels in 520 human Head and Neck Squamous Cell Carcinoma (HNSCC) tumor tissues obtained from TCGA. (G) FARDEEP deconvolution tool was used to categorize immune cell subsets. High SLC2A1 expression was found to be inversely correlated with frequencies of CD8+ cytotoxic T lymphocytes. Memory T cells, follicular helper T cells, and M1-like macrophages. Conversely, *SLC2A1* expression was moderately positively correlated with the frequency of regulator T-cells (Treg).

Hypoxia is a key feature of HNSCC, which shows a median partial pressure of oxygen (pO_2_) of 9 mmHg, in contrast with a pO_2_ of 40-60 mmHg in the paired normal tissue^6^. To assess whether the Sox2-GEMM recapitulates this characteristic metabolic alteration, we first dissected buccal mucosa from homozygous Sox2-GEMM and K5-CreER control mice five weeks post-tamoxifen oral painting. We found that the expression levels of *Hif1a* and *Slc2a1* were significantly increased in transformed buccal mucosa (Figure 2E and S1A). Slc2a1 encodes a glucose transporter to support the high glycolysis demand in cancer cells and myeloid cells while T-cells express high levels of *Slc2a3*, another glucose transporter, in the TME^7^. A meta-analysis of 37 studies involving 3,272 HNSCC patients showed that the expression levels of *SLC2A1* were inversely correlated with overall survival, disease-free survival, and recurrence-free survival^8^. We stained normal mucosa, transforming OED two weeks post-induction and HNSCC five weeks post-induction with a Slc2a1-specific antibody. In normal mucosa, the Slc2a1 stain was either negative or showed a weak cytoplasmic staining pattern. However, in transforming OED, Slc2a1 assumed a conspicuous membranous staining pattern in the lesional areas, with a clear demarcation from the adjacent and relatively normal epithelium, which lacked the membrane staining pattern. HNSCC showed a diffuse, strong membranous staining pattern (Figure 2E). Notably, the infiltrating lymphocytes were largely negative for Slc2a1 in both OED and HNSCC (Figure 2E). This finding is consistent with a TCGA analysis, in which we found that HNSCC expressed significantly higher levels of SLC2A1 than control mucosa (Figure 2F).

To explore the functional impact of high SLC2A1 expression in HNSCC on the tumor immune infiltrate, we deconvolved the immune landscape of 520 TCGA HNSCC patients, using a robust tool, FARDEEP^9, 10^. We found that the expression levels of *SLC2A1* were significantly inversely correlated with the frequencies of CD8^+^ CTL (ρ=-0.16, ***p=0.00021), memory T-cells (ρ=-0.13, **p=0.0029), follicular helper T-cells (ρ=-0.098, *p=0.026), γδ T-cells (ρ=-0.28, ****p<0.0001), and M1-like macrophages (ρ=-0.30, ****p<0.0001). The expression levels of *SLC2A1* were moderately positively correlated with the frequency of regulatory T-cells (Treg) (ρ=0.088, *p=0.046) (Figure 2G). Then, we separated the HPV^-^ and HPV^+^ HNSCC and performed immune deconvolution. We found that the overall impact of high SLC2A1 expression in the tumor immune microenvironment is similar between HPV^-^ and HPV^+^ HNSCC, with HPV^-^ tumors showing a more prominent correlation. We found that high levels of SLC2A1 were inversely correlated with CD8^+^ CTL (ρ=-0.13, **p=0.0091), γδ T-cells (ρ=-0.24, ****p<0.0001), and M1-like macrophages (ρ=-0.31, ****p<0.0001) in HPV^-^ tumors (Figure S2A-S2C).

### SOX2-driven OED transformation remodels immune infiltrates in longitudinal human specimens

To verify the TCGA analysis, we procured longitudinal specimens from patients with SOX2^+^ OED that had progressed into invasive HNSCC after excision of the pre-cancerous lesion. In normal oral mucosa, SOX2 stains sporadic cells at the basal layer with notable spacing. In this group of specimens, SOX2 staining was noted above the basal epithelial layer in the OED specimens. Upon malignant transformation, SOX2 showed a diffuse and strong nuclear staining pattern (Figure 3A). We performed multispectral immunofluorescence full-slide staining using a lymphoid cell panel and a myeloid cell panel. We selected 6-13 fields of interest per slide based on the presence of dysplasia, tumors, or dense immune infiltrates. We examined the lymphoid cell markers, including CD8, CD4, GATA3, PD1, CXCR3, and TBET as well as myeloid cell markers CD33, CD68, CD11c, HLR-DR, and PD-L1 (Figure 3B). We found that SOX2-driven transformation significantly reduced TBET^+^CD8^+^ T-cells and CXCR3^+^CD8^+^ T-cells (Figure 3C and 3D), two effector subsets that express high levels of IFN-γ^11, 12, 13, 14^. The transformation also resulted in a significant expansion of PD-L1^+^CD33^+^ myeloid cells (Figure 3E). Similarly, matched SOX2^+^ HNSCC contained significantly increased levels of PD-L1^+^CD68^+^ macrophages/monocytes and PD-L1^+^CD11c^+^ dendritic cells (Figure S3A and S3B). We also noted that there were more HLA-DR^+^ myeloid cells in SOX2^+^ HNSCC than matched OED (Figure S3C-3E).

**Figure 3.**
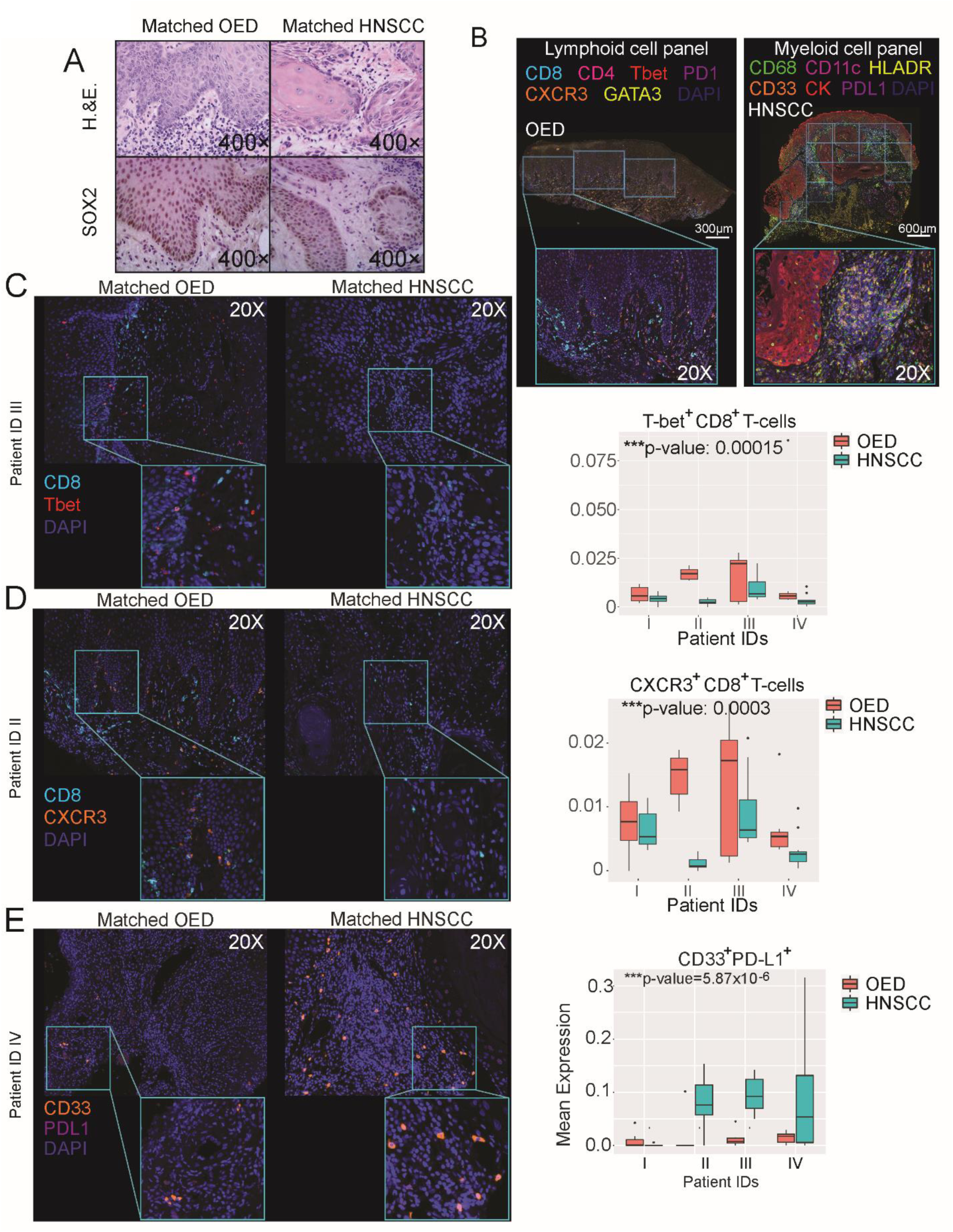
SOX2+ OED to HNSCC lesions exhibit a hypoimmunogenic tumor microenvironment. (A) H&E and immunohistochemical staining of paired SOX2^+^ oral epithelial dysplasia (OED) and head and neck squamous cell carcinoma (HNSCC) tissue specimen. (B) Representative multispectral immunofluorescence images (MSI) images of tumor infiltrating lymphoid and myeloid cells in paired SOX2^+^ OED/HNSCC patient specimen. Lymphoid panel MSI stains included the following: anti-CD8 (cyan), anti-CD4 (magenta), anti-Tbet (red), anti-PD1 (purple), anti-CXCR3 (orange), anti-GATA3 (yellow), and DAPI (blue). Myeloid panel m-IF stains included the following: anti-CD68 (green), anti-CD11c (magenta), anti-HLADR (yellow), anti-CD33 (orange), anti-cytokeratin (CK, red), anti-PDL1 (purple), and DAPI (blue). (C-E) MSI images were taken using the Akoya Biosciences’ Vectra Polaris scanner and regions of interest were selected using Akoya Phenochart software. We conducted tissue segmentation, cell segmentation, and cell phenotyping using Akoya inForm analysis software. Further data analysis was performed by first consolidating the data in Akoya phenoptrReports package in Rstudio and then performing a two-sample Wilcoxon test comparing the mean expression level between the OED and HNSCC stages for each patient. (C) Representative MSI images of CD8+Tbet+ immune infiltrate paired OED/HNSCC tissue specimen. (D) Representative MSI images of CD8+CXCR3+ immune infiltrate paired OED/HNSCC tissue specimen. (E) Representative MSI images CD33+PDL1+ immune infiltrate in paired OED/HNSCC tissue specimen. Results were considered significant if the p-value was less than 0.05.

### SOX2-driven OED transformation remodels the lesional immune landscape as a function of time

To better characterize the longitudinal lesional immune landscape as a function of time during malignant transformation, we induced the homozygous Sox2-GEMM with tamoxifen painting on the oral mucosa for five consecutive days. We then collected live CD45^+^ immune cells from bilateral buccal mucosa two weeks post-tamoxifen painting, during the OED stage, and four weeks post-induction when HNSCC showed 100% penetrance for histology-verified HNSCC development. Flow cytometric analysis revealed a significant reduction in both the overall CD4^+^ and CD8^+^ T-cells in cancer compared to mucosa-resident immune cells from K5-CreER control (Figure 4A and 4B). Previous studies using human HNSCC specimens show that PD-1^low^ T-cells showed stronger effector function in the TME and correlated with better outcomes^15^. Thus, we evaluated effector T-cells based on their expression levels of immune checkpoint receptors. We found that both CD4^+^PD-1^low^ and CD8^+^PD-1^low^ subsets were decreased upon oral mucosal malignant transformation (Figure 4C and 4D). Notably, the levels of CD4^+^Tim3^low^ and CD8^+^Tim3^low^ effectors were significantly excluded from Sox2-driven HNSCC compared to the normal oral mucosa (Figure 4E and 4F). OED assumed an intermediary phenotype for these markers.

**Figure 4.**
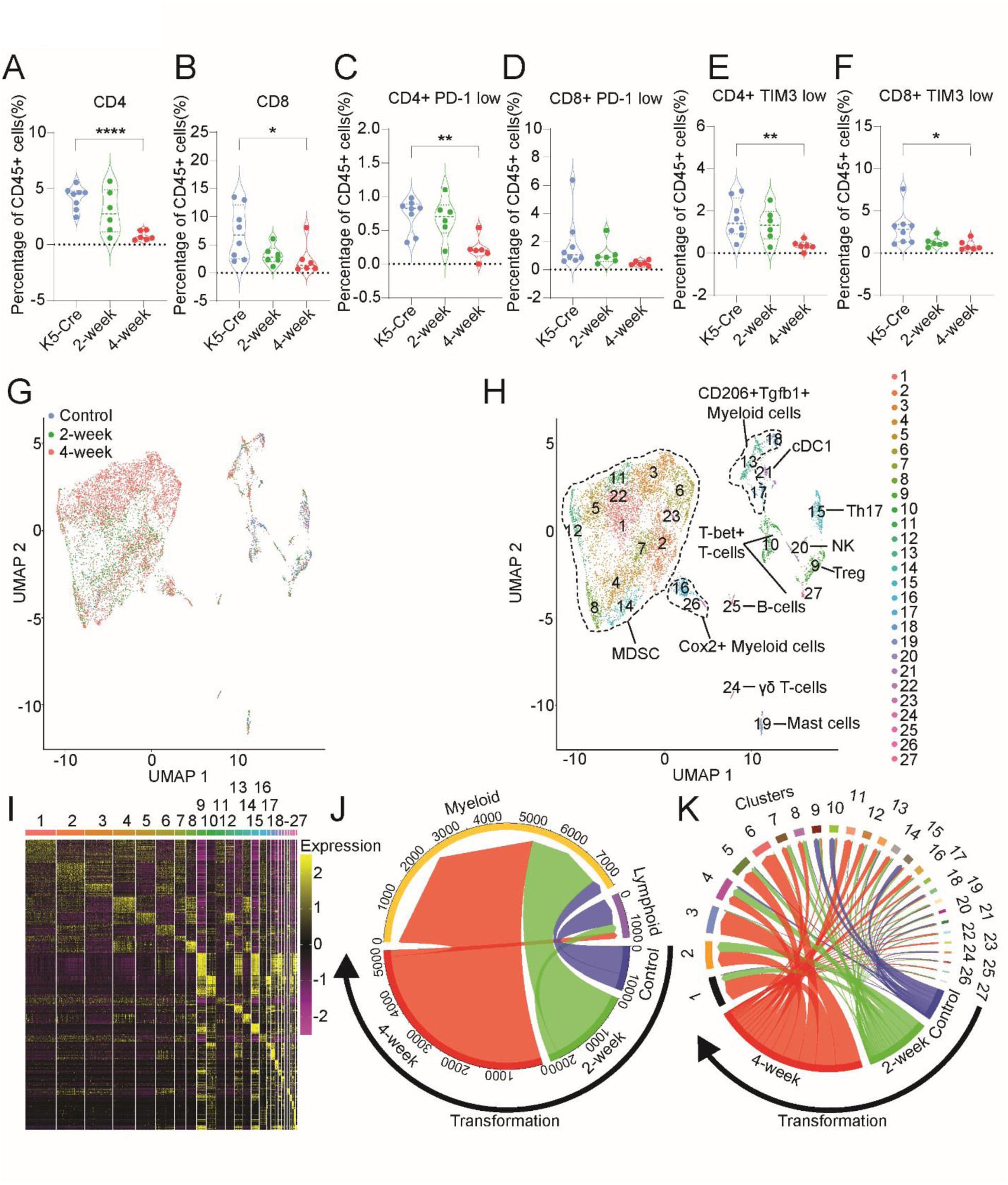
Major immune subsets in Sox2-mediated oral epithelial dysplasia and cancer. (A-F) Flow cytometric analysis on immune cell infiltrates in the buccal mucosa of *K5-CreER^+^;Sox2^+/+^* painted with tamoxifen to induce epithelial Sox2 expression for 2-weeks (n=6, oral epithelial dysplasia stage) and 4-weeks (n=6, oral cancer stage) compared to *K5-CreER^+^ control mice* (n=8). Representative violin plots show the percentage of expression for each marker indicated; and error bars represent mean ± SEM. Cells were gated from single/Zombie Aqua^-^ live, CD45^+^ TCRb^+^, and then further gated on CD4^+^ (A) and CD8^+^ (B) cells. Further gating from the CD4+ and CD8+ populations, we explored the surface level expression of PD-1 (C-D) and TIM3 (E-F). Comparisons between the three groups were made using a one-way ANOVA (*p<0.05; **p<0.01; ****p<0.0001). (G-H) Integrated UMAP of immune cell populations in the buccal mucosa of tamoxifen-painted *K5-CreER^+^;Sox2^+/+^* mice, 2- and 4-weeks post induction, and *K5-CreER^+^* mice. Buccal mucosal tissue was harvested and CD45+ tumor-infiltrating immune cells were FACS sorted and subjected to single-cell RNA sequencing. (I) Heatmap showing the average expression of the top differentially expressed genes for the major cell types identified in buccal mucosal tissue. (J-K) Chord diagram depicting the abundance of myeloid and lymphoid cells in control (blue), 2-weeks (green), and 4-weeks (red). Further analysis of the chord diagram was performed to show the abundance of the 27 identified cell clusters in each group are shown.

The flow cytometric immune profiling showed a T-cell exclusion phenotype in established HNSCC, also seen in clinical specimens, lending further support to the rigor of this model. However, it was still unclear how the TME evolved into this terminal stage given the somewhat intermediary T-cell phenotype in OED. Thus, we sought to increase our immune phenotyping resolution by using single-cell (sc)RNA-Seq. After rigorous filtering, we obtained 8,960 high-quality transcriptomes during oral mucosa malignant transformation. We previously showed that including a panel of lineage marker genes in addition to the most differentially expressed genes in immune single-cell data sets stabilized immune subset segregation and improved the rigor of annotation^16, 17^. We employed a similar method to render divergent immune lineages. In addition, we performed Single-cell Analysis Via Expression Recovery (SAVER) imputation to improve immune signature gene expression profiling. Epithelial expression of Sox2 drove a massive expansion of inflammatory cells, which clustered mostly in the left side of the UMAP (−10 to 5 UMAP X-axis region) (Figure 4G). We identified 27 clusters with distinct gene expression patterns. This overall examination revealed that most of the inflammatory cells that emerged in the OED and HNSCC stage were suppressive myeloid cells, such as MDSC (clusters 1, 2, 3, 4, 5, 6, 7, 8, 11, 12, 14, 22, and 23) and Ptgs2^+^ myeloid cells (clusters 16 and 26) (Figure 4H and 4I). Overall, Sox2-driven epithelial malignant transformation entailed a massive expansion of myeloid cells alongside a concurrent loss in lymphocytes (Figure 4J). Notably, most T-cells identified across time points of HNSCC evolution were from the healthy control oral mucosa group (Figure 4J and 4K). A closer assessment showed that nearly all MDSC were only identifiable in the OED and HNSCC stage (clusters 1, 2, 3, 4, 5, 6, 7, 8, 11, 12, 14, 22, and 23) (Figure 4J and 4K). In contrast, normal control oral mucosa contributed the majority of Batf3^+^ cDC1 conventional dendritic cells type 1 (cDC1, cluster 21), Tbet^+^ T-cells (clusters 10 and 27), NK cells (cluster 20), and γδ T-cells (cluster 24) (Figure 4K).

The above analysis allows us to compare similar immune subsets across time points. However, certain functional states may arise de novo during the transformation process and are not identifiable in normal mucosa. To identify unique immune functional states emerging at the early transformation stage of the oral mucosa, we next individually analyzed the immune subsets at the three stages of the disease spectrum. Normal oral mucosa contained balanced CD8^+^ T-cells (cluster 17), NK cells (cluster 3), NKT cells (cluster 14), γδ T-cells (clusters 6 and 10), Treg (cluster 1), Lag3^+^ T-cells (cluster 16), Th2 cells (clusters 4 and 8), and Th17 cells (cluster 5). The myeloid cells contained subsets expressing high levels of *Tnf* (clusters 13), *Batf3* (cluster 12), and small subsets expressing *Tgfb1* (clusters 7 and 9). The overall architecture of oral mucosa-resident immune cells had a high degree of heterogeneity with balances and checks (Figure 5A). After we induced oral epithelial expression of Sox2, the lesion started to progress to the OED stage. At this time point, no invasive HNSCC histology was identified. However, the immune landscape underwent dramatic shifts with an apparent loss of population heterogeneity. NK cells, NKT cells, and γδ T-cells were the first to disappear at this very early stage of disease progression. In contrast, an expansive group of Tcr^-^Ly6c^-^Ly6g^+^ MDSC emerged with a subset of them expressing high levels of PD-L1 (clusters 1, 3, 5, 7, and 10) and the other subset being largely PD-L1^-^ (clusters 2 and 4) (Figure 5B). As OED progressed into invasive HNSCC, CD8^+^ T-cells were largely lost at this time, in agreement with our flow cytometric findings and those seen in clinical specimens. Treg and Th17 cells persisted in these advanced lesions. The most striking change was that myeloid cells expanded substantially with progressively complex immune suppressive features. Clusters 1, 2, and 3 expressed high levels of *Nos2*, which sustains MDSC fate and function^18, 19^. Clusters 4, 6, and 9 were MDSC subsets that expressed high levels of PD-L1 and *Il10*. Clusters 5, 7, 8, and 11 form another subset of Ly6g^+^ MDSC that expressed high levels of Il1b and variable levels of *Ptgs2*, *PD-L1*, and *Lgals9*. Clusters 10, 12, and 13 were myeloid cells distinguished by high expression levels of *Tgfb1* (Figure 5C). Overall, at this stage of the disease, the myeloid cells had evolved into subsets with compensatory immune suppressive features. Some cells were PD-L1^+^ and many were PD-L1^-^ but featured other suppressive features.

**Figure 5.**
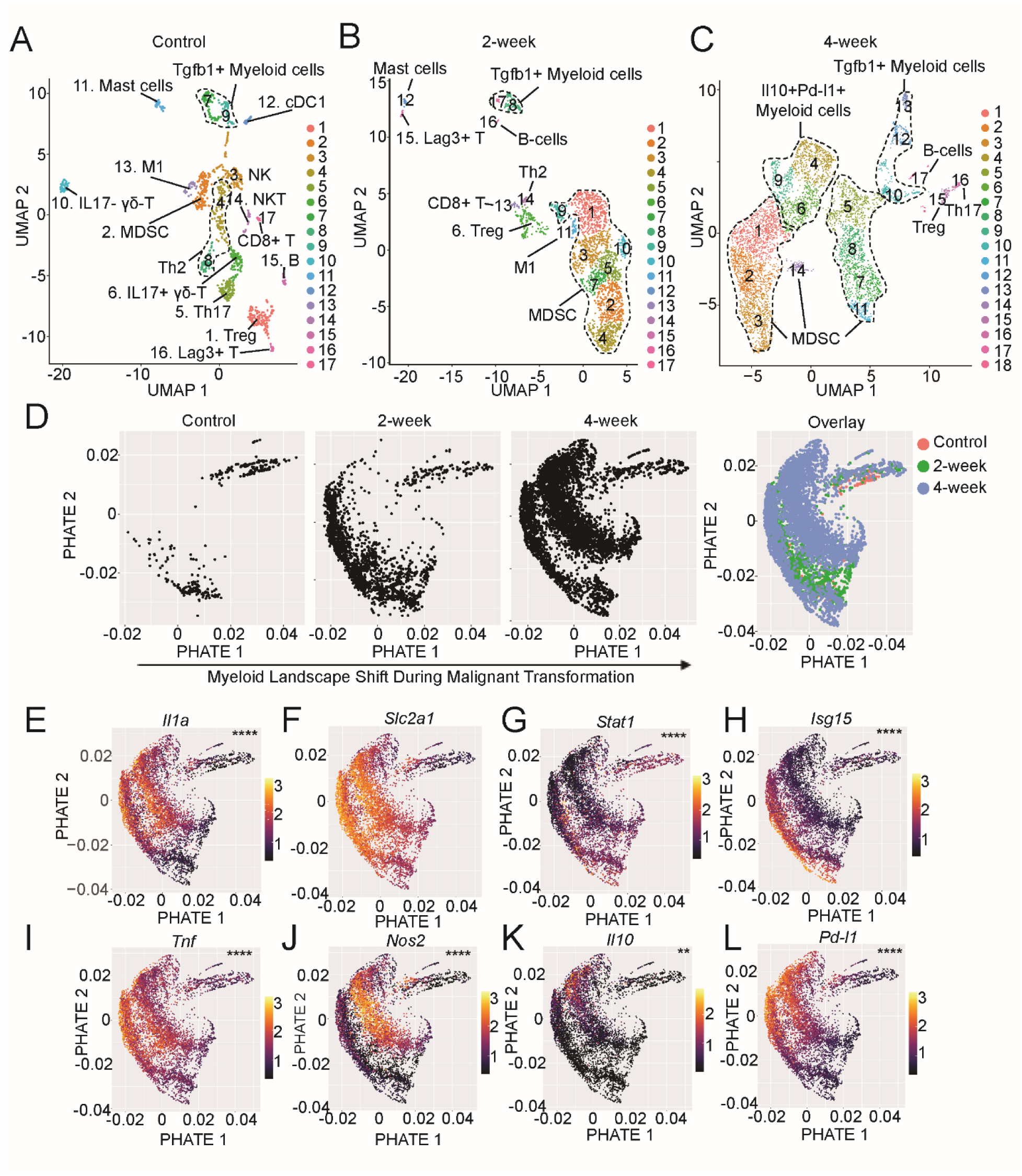
Expansion of hypoimmunogenic myeloid cells following Sox2-mediated malignant transformation. (A-C) Representative UMAP of the major immune cell types identified by single-cell RNA sequencing of buccal mucosal tissues from *K5-CreER^+^;Sox2^+/+^*mice at 2-weeks and 4-weeks post tamoxifen administration, and K5-CreER+ control mice. (D) Integrated PHATE mapping of myeloid cells at 2-weeks and 4-weeks post tamoxifen administration, and in control mice treated with tamoxifen for 4 weeks. (E-L) Representative PHATE maps comparing the expression of genes in myeloid cells at 2-week and 4-week transforming lesions compared to the control. Myeloid cells from transforming lesions exhibited a dichotomous signature of high *Il1a, Slc2a1, Tnf,* and *Nos2* expression but low type I interferon activation as indicated by low *Stat1, Isg15 and Il10* expression.

### Myeloid cells in initiating Sox2-driven HNSCC assumed unique IL1^high^IFN^low^ signatures

Because the most dramatic immune subset changes in Sox2-driven HNSCC were centered around myeloid cells, we separated the myeloid cells and performed the potential of heat diffusion for affinity-based transition embedding (PHATE) analysis, which offers a powerful visualization tool to identify cellular functional differentiation trajectories^20^. Resident myeloid cells in healthy oral mucosa were primarily present in the lower left and upper right quadrants of the PHATE map. As soon as Sox2 was activated, myeloid cells started to expand in the local environment. Such expansion was likely a combination of in situ proliferation, shown in cell clusters continuous with those from healthy oral mucosa, and de novo recruitment, which was illustrated in clearly separated clusters on the PHATE maps (Figure 5D). We compared the gene expression patterns between the myeloid cells from transforming lesions at the two-week and four-week time points and those from normal oral mucosa. We found that the myeloid cells in transforming lesions assumed a distinct signature with high levels of *Il1a*, *Il1b*, *Il1rap*, *Slc2a1*, and *Hif1a*, but low levels of IFN-I signatures such as *Stat1*, *Isg15*, and Major Histocompatibility Complex (MHC) class II (Figure 5E-5H, S4A-S4C, and Table S1). The myeloid cells from transforming lesions expressed high levels of *Tnf*, *Nos2*, Il10, and Pd-l1 (Figure 5I-5L). Then, we performed gene set enrichment analysis (GSEA) between normal mucosa-resident and intra-lesional myeloid cells and found that the most significantly altered pathways included defense response, cell death, response to stress, amide metabolism, cell migration, cytokine production, and innate immune response (Figure S4D). We also characterized the most significantly altered signaling pathways, which included mitogen-activated protein kinases (MAPK), Il6 signaling, Tnf signaling, small GTPase signaling, NK-κB signaling, glycolysis pathway, TLR signaling, pyruvate metabolism, and response to IFN-I (Figure S4E).

### IL1α priming desensitized myeloid cell response to IFN-I agonists

To understand how Sox2 in tumor cells drove the massive expansion of lesion-infiltrating myeloid cells, we first employed an implantable HNSCC model. We found that Sox2-expressing tumors grew more aggressively, consistent with our previous findings (Figure 6A). Sox2-expressing HNSCC contained significantly fewer CD8^+^ T-cells and CD8α^+^ cDC1 in the TME (Figure 6B and 6C). In contrast, Sox2^+^ HNSCC contained significantly elevated levels of MDSC (Figure 6D). To determine how Sox2^+^ HNSCC created this more suppressive MDSC-rich TME, we performed RNA-Seq analysis of empty vector control and Sox2-expressing tumor cells. Among the most significantly changed genes were *Il1a*, *Csf2,* and *Ccl2* (Figure S5A, Table S2). Because CCL2 is a principal chemotactic agent for myeloid cell recruitment, we validated findings in four murine HNSCC cell lines, including MOC1, MOC2-E6/E7, NOOC1, and NOOC2. Sox2 expression consistently upregulated the production of Ccl2 (Figure 6E-H). Then, we further verified findings in human HNSCC cell lines UMSCC22A and UMSCC108 and confirmed the SOX2-CCL2 axis (Figure 6I-J). Thus, SOX2 amplification in transforming epithelial cells promotes a potent chemotactic signal, CCL2, to promote the recruitment of myeloid cells featuring a high IL1, high glycolysis, and low IFN transcriptional program.

**Figure 6.**
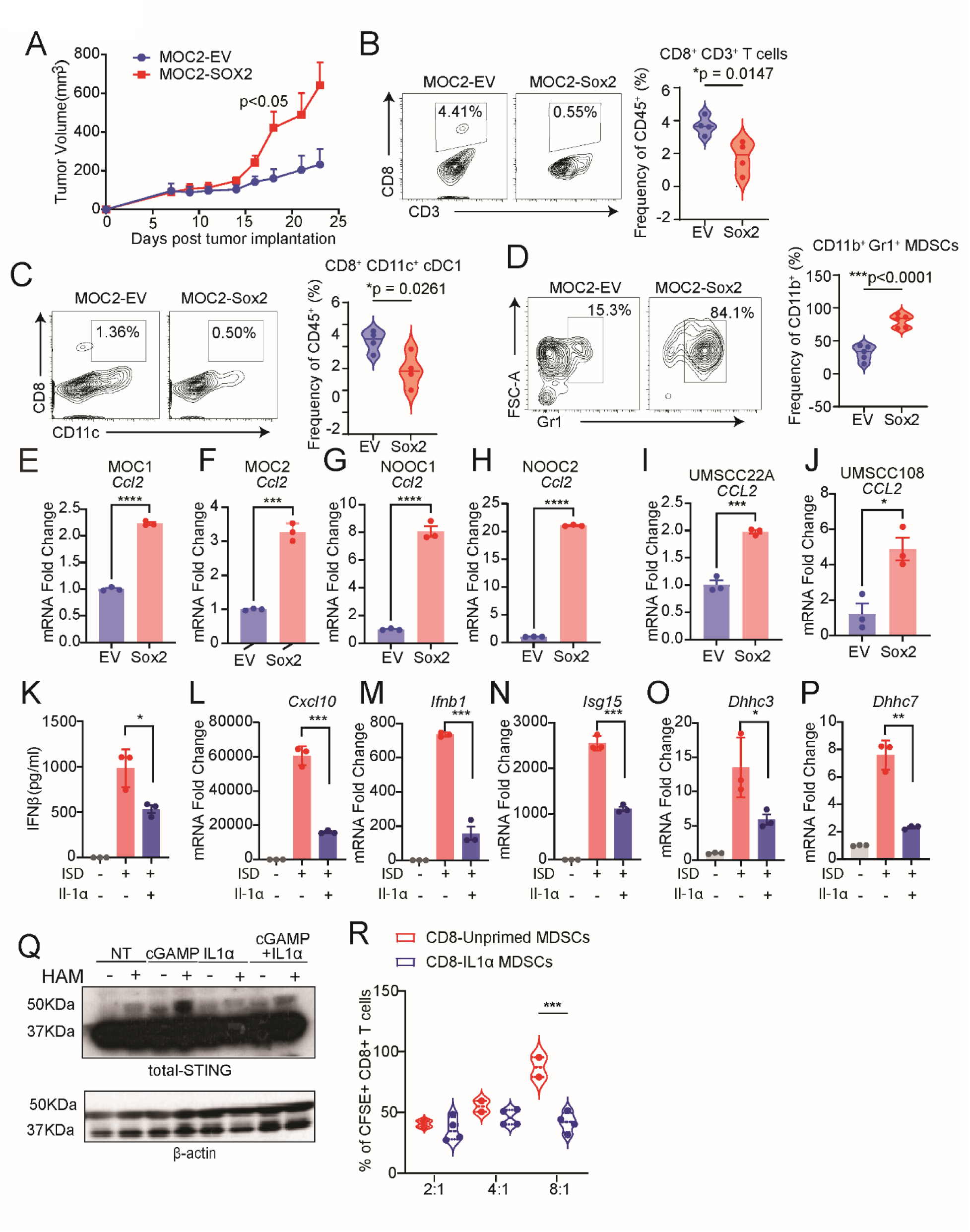
Impact of IL1α priming on tumor growth and myeloid cell STING responsiveness and function. (A) One million MOC2-Sox2 or MOC2-EV oral cancer cells were implanted subcutaneously into the right flank of C57BL/6J mice. Tumor growth was monitored every other day starting on day 7 post-implantation. Results represent mean ± SEM, and the growth curves between the two groups was compared using GEE. (B-D) Representative flow cytometry scatterplots and violin plots from tumor homogenates Ficoll-separated immune cells from MOC2-Sox2 (n=4) and MOC2-EV (n=4) tumors. Cells were gated from single/Zombie Aqua-live, CD45^+^ cells, and then further gated into TCRb^+^ CD8^+^ CD3^+^ T cells (B) and CD8^+^ CD11c^+^ conventional dendritic cells (cDC1) (D). (E-J) Total RNA was isolated from murine MOC1, MOC2, NOOC1 and NOOC2, human UMSCC22A and UMSCC108 HNSCC cell lines expressing either *EV* or *SOX2* to quantify the mRNA expression levels of *CCL2.* Values represent mean ± SEM of three technical replicated per sample. Comparisons were made using t-test (*p<0.05, ***p<0.001, ****p<0.0001). (K) Bone marrow-derived macrophages (BMDMs) were primed with 10ng/mL recombinant murine proteins IL1α for 24 hours, rested for 4 days, and then challenged with a STING agonist (2µ/mL ISD) for 16 hours. Supernatant was collected and assessed for Ifn-β by ELISA. Values represent mean ± SEM of three technical replicated per sample. Comparisons were made using t-test (*p<0.05, **p<0.01, ***p<0.001). (L-P) Total RNA was isolated from IL1α-primed BMDM challenged with ISD to quantify the mRNA expression levels of *Cxcl10, Ifnb1, Isg15, Dhhc3,* and *Dhhc7.* (Q) To assess the impact of IL1α priming on STING palmitoylation, we treated Thp1 monocytes with recombinant human IL1α protein for 24 hours, rested for 4 days, and then challenged with 10µg/mL cGAMP for 30 minutes. Protein lysates were isolated and subjected to an Acyl-PEG exchange assay, which uses cysteine-specific click chemistry to exchange S-fatty acylation sites with a mass tag of a defined size (5kDa, methoxypolyethylene glycol maleimide), observed by total-STING immunoblot. This assay cleaves thioester bonds coupling fatty acid residue to the protein using hydroxylamine (NH_2_OH), with the absence of NH_2_OH serving as an internal control. Palmitoylated STING protein appears 10kDa above its known molecular weight (35kDa). (R) To assess the functional impact of IL1α priming on the myeloid derived suppressor cells (MDSCs), we performed a T cell suppression assay. Bone marrow-derived MDSCs were either primed with 10ng/mL IL1α or left unprimed, rested for 4 days, activated with 0.1µg/mL LPS and 100 U/mL IFN-γ for 16 hours to become activated-MDSCs. CFSE-labeled CD8^+^ T cells were cultured with IL1α-primed and unprimed MDSC for four days at CD8^+^ T cells: MDSC ratios of 2:1, 4:1, and 8:1, and assessed by flow cytometry for CFSE dilution. Volin plots show the percentage of CFSE^+^ CD8^+^ T cells, with data representing two biological replicates in the unprimed MDSC group and four biological replicates in the IL1α-primed MDSC group. Comparisons between the two groups were made using a two-tailed, unpaired t-test (***p<0.001).

Since we identified a distinct IL1α ^high^IFN^low^ signature for the lesional myeloid cells during the malignant transformation of epithelium, we sought to investigate whether IL1α signaling was either a correlational or causative factor impacting myeloid cell plasticity in IFN response. We pulsed bone-marrow-derived macrophages (BMDM) with a low dose of Il1α (10 ng/mL) for 16 hours and then washed the cells with PBS. The BMDM were rested for four days before we challenged them with STING agonists ISD and cGAMP. We found that the brief Il1α priming desensitized BMDM response to STING agonists, with significantly lower expression levels of IFN-I signature genes and protein levels of IFN-β in the supernatant (Figure 6K-N). The activation of STING requires its palmitoylation, the addition of fatty acid residues to free cysteine domains, which facilitates ER-to-Golgi translocation and recruitment of effector molecules^21^. We found that IL1α priming reduced the expression of both *Dhhc3* and *Dhhc7,* which encode the palmitoyltransferases responsible for the palmitoylation of STING (Figure 6O-P). Then, we employed click chemistry to assess whether Il1α priming in THP-1 cells reduced total palmitoylated STING. Exchanging palmitoylated fatty acid residues on STING with acyl-PEG followed by immunoblotting revealed that IL1α priming inhibited the palmitoylation of STING (Figure 6Q). To directly assess the functional impact of IL1α priming on the MDSCs, we cocultured Il1α-primed MDSCs with CFSE-labeled CD8^+^ T cells for four days at different ratios. The CFSE^+^CD8^+^ T-cells were quantified by flow cytometry (Figure S5B and S5C). Compared with unprimed MDSCs, Il1α-primed BMDCs inhibited the proliferation of T-cells more potently (Figure 6R). Collectively, we identified a mechanistic link between early IL1α exposure and suppressive myeloid cell differentiation during epithelial malignant transformation.

### Blockade of early IL-1 signaling delays the onset of Sox2-driven HNSCC

To assess the pathological relevance of IL1 signaling in epithelial transformation, we first treated induced *K5-CreER;Sox2^+/+^*mice with 200 μg anti-Il1r1 intraperitoneal (i.p.) injections. These injections were given concurrently with tamoxifen application on days 0 and 3 and were followed by additional injections after induction on days 6 and 9. Il1r1 blockade significantly extended the survival of these mice (Figure 7A). In agreement, anti-Il1r1 resulted in a notable decrease in PD-L1^+^ MDSCs within the progressing buccal lesions (Figure 7B). Then, we crossed the *K5-CreER;Sox2^+/+^* GEMM with *Il1r1^-/-^* mice to generate the *K5-CreER;Sox2^+/+^;Il1r1^-/-^*compound strain. Consistent with the pharmacological approach in antagonizing the IL1 pathway, deletion of *Il1r1* from the *K5-CreER;Sox2^+/+^*strain significantly improved overall survival, nearly doubling median survival (Figure 7C). Because of the rapid tumor onset in this model, most buccal cancers were microscopic, we compared the areas where invasive tumor islands were identified and found that *K5-CreER;Sox2^+/+^;Il1r1^-/-^* mice showed significantly reduced areas of tumor than *K5-CreER;Sox2^+/+^;Il1r1^+/+^*littermate controls (Figure 7D). We extracted lesion-infiltrating immune cells and performed flow cytometry. We found that deletion of *Il1r1* significantly reduced the infiltrating Glut1^+^CD206^+^CD11b^+^ myeloid cells (Figure 7E). To profile the TME, we harvested buccal mucosa from both groups and performed RNA-Seq. The most significantly upregulated pathways in the *K5-CreER;Sox2^+/+^;Il1r1^-/-^* group compared with the littermate control *K5-CreER;Sox2^+/+^;Il1r1^+/+^* mice were IFN-γ response, allograft rejection, and IFN-α response, suggestive of enhanced Th1 immunity (Figure 7F).

**Figure 7.**
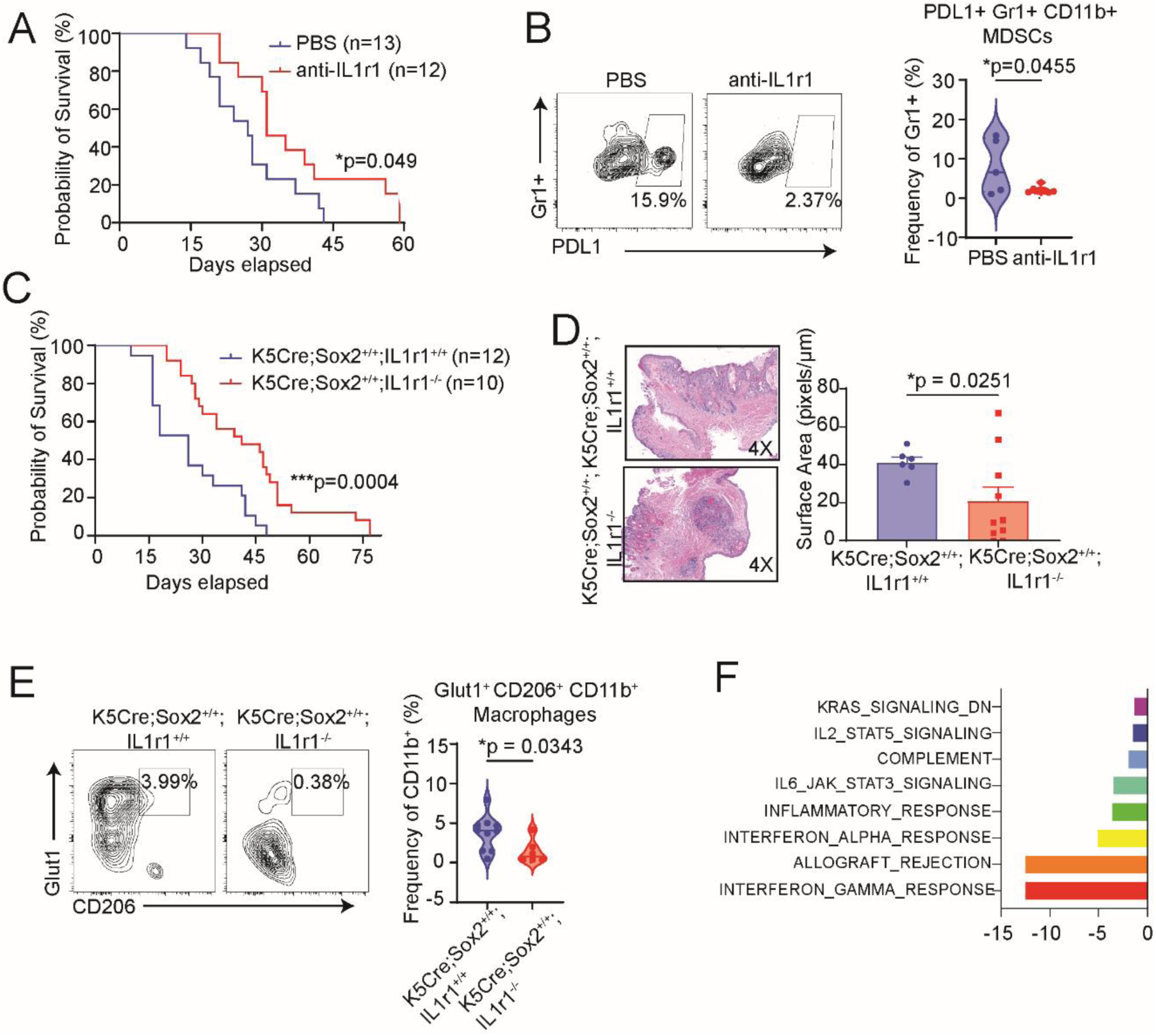
Blockade of IL1/IL1r1 signaling delays onset of Sox2-driven oral cancer and modifies innate immune response.\. (A) Longitudinal survival study in tamoxifen-painted *K5-CreER^+^;Sox2^+/+^*mice treated with 2mg/mL anti-IL1r1 (n=13) or PBS (n=12) via intraperitoneal injection on days 0, 3, 6, and 9. Body weights were monitored from the first day of tamoxifen application until body weight dropped below 20% of the initial body weight. The study was repeated twice, and differences in body weight between groups were assessed using the generalized estimating equation (GEE) model (*p<0.05). (B) Buccal mucosal tissue was harvested from *K5-CreER^+^;Sox2^+^* mice treated with either anti-IL1r1 or PBS and subjected to flow cytometric analysis. Representative scatterplots and violin plots show the percentage of expression of PDL1^+^ MDSCs. Cells were gated from single/Zombie Aqua-live, CD45^+^, CD11b^+^, and then further gated on Gr1^+^ (Ly6g/c) and PDL1^+^ cells. Comparisons between the two groups was performed using a two-tailed unpaired t-test (*p<0.05) (C) Longitudinal survival study in tamoxifen-painted *K5CreSox2^+/+^;IL1r1^+/+^*(n=12) and *K5CreSox2^+/+^;IL1r1^-/-^* (n=10) mice. Body weights were monitored from the first day of tamoxifen application until a drop in 20% initial body weight was observed. The study was repeated twice, and differences in body weight between groups were assessed using the GEE model. (D) Buccal mucosal tissue was harvested from *K5CreSox2^+/+^;IL1r1^+/+^* (n=6) and *K5CreSox2^+/+^;IL1r1^-/-^* (n=10) mice. Representative H&E staining patterns are shown. Surface area quantification of tumors was determined using ImageJ. Values represent mean ± SEM and the two groups were compared using a two-tailed unpaired t-test (*p<0.05). (E) Representative scatterplots and violin plots show the percentage of Glut1^+^ CD206^+^ CD11b^+^ Macrophages. Cells were gated from single/Zombie Aqua-live, CD45^+^, and then further gated on CD11B^+^CD206^+^ Glut1^+^ cells. Comparisons between the two groups were made using a two-tailed unpaired t-test (*p<0.05). (F) Buccal mucosal tissue from *K5CreSox2^+/+^;IL1r1^+/+^* (n=5) and *K5CreSox2+/+;IL1r1-/-* (n=5) were harvested and subjected to bulk RNA sequencing. A bar graph of top pathways altered between groups is shown.

## Discussion

HNSCC has a modest, usually below 15%, response rate to ICI. Once HNSCC is established, effector T-cells assume terminal exhaustion phenotypes. Even targeting multiple T-cell subsets with a combination of anti-PD-1 and anti-CTLA4 did not show a clinical benefit over a single agent alone^22^. Thus, the identification of high-risk pre-cancerous lesions and a better understanding of the early-stage immune alteration hold key promise to reducing HNSCC burden.

Implantable HNSCC models do not recapitulate many histologic features essential to render the diagnosis of squamous cell carcinoma. More importantly, the establishment of implantable models does not undergo a precancerous phase, which makes it less helpful to understand the early epithelial-immune interactions during malignant transformation. This study integrates both longitudinal human specimens and a unique GEMM to show that SOX2^high^ OED bears a heightened risk for progressing into hypoimmunogenic oral cancers. We characterized a signaling axis that starts with SOX2-induced CCL2 expression in transforming epithelial cells, which recruits myeloid cells with significantly increased IL-1α and glycolysis signals and dampened IFN-I signatures. Notably, the remodeling of the myeloid cell compartment occurs before the emergence of the histology of invasive HNSCC, making it a fate-committing early event.

Previous studies have highlighted the roles of IL-1α in tumor progression in breast cancer^23^, liver cancer^24^, gastric cancer^25^, and lung cancer^26^. IL-1α can promote the proliferation and survival of HNSCC cells in an NF-κB- and AP-1-dependent fashion and downregulate CD80 costimulatory molecule^27, 28^. An analysis of 154 HNSCC patients found that IL-1α strongly correlates with distant metastasis^29^. Tumor-derived IL-1α can stimulate the expression of an alarmin family member, thymic stromal lymphopoietin, which promotes tumor cell survival^23^. However, how IL-1α modulates the anti-tumor immune response in precancerous lesions was unclear. This study identified a link underpinning myeloid cell desensitization to IFN-I agonists. We found that IL-1α priming potently inhibits the palmitoylation of STING, leading to the desensitization to IFN-I agonists and a more suppressive phenotype in a subset of myeloid cells. Using pharmacological and genetic approaches, we found that early blockade of IL1 signaling reduces the infiltration of subsets of highly suppressive myeloid cell subsets and delays the onset of Sox2-driven HNSCC. Many STING-inducing strategies, such as irradiation, inhibition of DNA damage repair, and delivery of STING agonists, are being evaluated in clinical trials to sensitize cancers to ICI. Our results suggest that myeloid cells that experienced IL-1α in the TME are insensitive to STING stimulation. IL-1α-expressing myeloid cells are a dominant population during early epithelial transformation. Future studies are needed to explore strategies to resensitize these IL-1α-experienced myeloid cells to innate immune agonists.

Overall, we integrated longitudinal human specimens and mouse models to show that high SOX2 expression in epithelial cells is a high-risk event, engendering a SOX2-CCL2 pathway between transforming epithelial cells and myeloid cells, the latter of which exhibits a distinct IL1^high^Glycolysis^high^IFN-I^low^ signature. As a mechanism, we found that IL-1α priming epigenetically reduces the expression of DHHC to inhibit the palmitoylation of STING, expanding MDSC. The WHO histologic grading carries little prognostic value for OED^30, 31^. Surgical resection cannot reverse field cancerization in a key subset of OED that progresses despite vigilant monitoring. Our findings help identify a subset of OED with true high risks and reveal potential early intervention points to improve HNSCC prevention.

## Methods

### Animal Studies

All animal handling and procedures were conducted in compliance with protocols approved by the Institutional Animal Care and Use Committee (IACUC) at the M.D. Anderson Cancer Center (00002478-RN00) and the University of Michigan (PRO00010232). Female *ROSA26-CAG-Sox2-GFP* mice were generously provided by Dr. Jianwen Que at Columbia University and crossed to male *K5-CreER* mice (RRID:IMSR_JAX:018394), which were purchased from the Jackson Laboratory. *Il1r1^-/-^* mice were purchased from the Jackson Laboratory (RRID:IMSR_JAX:003245) and crossed to *K5CreER;Sox2^+/+^* mice to generate *K5-CreER;Sox2^+/+^;Il1r1^-/-^* mice. All mice were group housed under specific pathogen-free conditions. To induce Sox2-mediated oral cancer, tamoxifen was applied topically to the oral mucosa for five consecutive days at a concentration of 40 mg/mL (T5648, Sigma-Aldrich). Each mouse was given 1 mg of tamoxifen dissolved in 25 μL of corn oil (C8267, Sigma-Aldrich) and applied while mice were anesthetized in an isoflurane induction chamber. In a subset of mice, we topically applied 5 mg/mL (25 μg dissolved in 20 μL of ethanol) DMBA three times within the same week when topical tamoxifen was applied. Both tamoxifen and DMBA were kept as stock solutions in a freezer and freshly prepared at the start of the treatment. To ensure that tamoxifen was completely dissolved in corn oil, 40 mg/mL solutions were incubated in a shaker at 37°C for two days.

### Cell culture

The HNSCC cell lines UMSCC108 and UMSCC22A were grown in DMEM (10-013-CV, Corning) with 10% fetal bovine serum (FBS) (Gibco, Life Technologies) and 100 U/mL penicillin-streptomycin (15-140-122, Gibco). THP-1 cells were cultured in RPMI 1640 (MT10040CV, Corning) containing 10% heat-inactivated FBS and the same concentration of penicillin-streptomycin. The MOC1, MOC2-E6/E7, NOOC1, and NOOC2 lines were maintained in a mixture of 60% IMDM (SH30228.01, HyClone) and 30% F12 nutrient mix (11764-054, Gibco) with 5% FBS, supplemented with 4 μg/mL puromycin, 5 μg/mL insulin, 40 ng/mL hydrocortisone, 5 ng/mL EGF, and 100 U/mL penicillin-streptomycin.

### Histology and Immunohistochemistry

Buccal mucosal tissue was harvested from mice, fixed using 4% paraformaldehyde overnight, and then transferred into 70% ethanol prior to paraffin embedding. Antigen unmasking was achieved by placing slides in citrate retrieval buffer (HK08120K, Biogenex) that was heated to a rolling boil for 10 minutes and then placed at room temperature for an additional 15 minutes before transferring slides to distilled water for a brief wash. Two subsequent washes in PBS-T were performed prior to placing slides in 3% hydrogen peroxide to quench endogenous peroxidase. After two PBS-T washes, tissue sections were incubated with 5% goat serum in PBS-T for one hour and then incubated at 4°C overnight with the following antibodies: SOX2 (23064, Cell Signaling Technology), Ki67 (ab16667, Abcam), p63 (39692S, Cell Signaling Technology), p53 (ab1431, Abcam), and SLC2A1/GLUT1 (20701, BiCell Scientific). After three washes in PBS-T, slides were incubated with an HRP-conjugated anti-rabbit secondary antibody (ab97051, Abcam) for 30 minutes at room temperature. We used an ABC kit (Cat. PK-4001, Vector Laboratories) for the detection of avidin-biotinylated HRP complexes. Counterstaining was performed using hematoxylin (NC9520196, Fisher Scientific), with the slides being immersed in the stain for 6 minutes. Slides were washed for 5 minutes in water and then dehydrated using ethanol at increasing concentrations and then xylene. Permount (SP15-100, Fisher Chemical) was used to mount the coverslip to the slides and then left to dry overnight. Slides were imaged and analyzed using ImageJ.

### Multispectral Imaging and Analysis

The lymphoid panel included the following markers: CD4 Opal 650 (1:50), CXCR3 Opal 540 (1:200), PD1 Opal 620 (1:200), CD8 Opal 690 (1:200), Tbet Opal 570 (1:50), Gata3 Opal 520 (1:100), and DAPI. The myeloid panel included the following markers: CD33 Opal 540, PDL1 Opal 620, CD11c Opal 650 (1:50), HLADR Opal 520 (1:250), CD68 Opal 690 (1:500), Cytokeratin Opal 570 (1:250), and DAPI. Images were taken using Akoya Biosciences’ (Akoya) Vectra Polaris scanner and regions of interest (ROI) were selected by a trained oral pathologist using Akoya Phenochart software. Spectral unmixing and image analysis were performed using Akoya InForm software, which was employed to develop and train algorithms for tissue segmentation, cellular segmentation, and phenotyping of tissue slides, enabling the classification of cell phenotypes for all markers of interest. Tissue segmentation was achieved by distinguishing tumor from stroma using cytokeratin staining, as well as identifying other regions such as glass. For cell segmentation and phenotyping, the ROI image data was converted into objects by identifying single cells based on their nuclear stain (DAPI), cytoplasmic, and membrane markers. Annotated files were imported into Rstudio v4.2.1. Akoyas’ R package phenoptrReports, an open-source tool that enables the consolidation of cell segmentation files generated in inForm. These consolidated data files contained slide summaries, cell counts and percentages, cell densities, mean expressions, H-scores, and nearest neighbor results. These datasets were further interpreted by first collapsing the metric of interest across multiple ROIs by the sample ID and then conducted a two-sample Wilcoxon test for every marker of interest. Mean expression level was compared between clinical stage (oral epithelial dysplasia vs head and neck cancer) for each patient. The test results are significant if the p-value is smaller than 0.05.

### Gene expression qPCR and ELISA

RNA isolation for qPCR was achieved using the QIAshredder (79654, Qiagen) and RNeasy Plus Mini Kit (74134, Qiagen). RNA concentration was measured by using a Nanodrop Spectrophotometer (Thermo Fisher Scientific). Reverse transcription of RNA into cDNA was achieved using a High-Capacity cDNA Reverse Transcription Kit and RNAse inhibitor (4368814 and N8080119, Applied Biosystems). The primers used for gene amplification were obtained from Integrated DNA Technologies, with the following specifications for each gene: *Sox2* F 5’- CCCACCTACAGCATGTCCTACTC, R 5’- TGGAGTGGGAGGAAGAGGTAAC; *Slc2a1* F5’- AGCCCTGCTACAGTGTAT,R 5’- AGGTCTCGGGTCACATC; *Cxcl10* F 5’- AATGAGGGCCATAGGGAAGC, R 5’- AGCCATCCACTGGGTAAAGG; *Isg15* F 5’- TGGAAAGGGTAAGACCGTCCT,R 5’- GGTGTCCGTGACTAACTCCAT; *Ifnb1* F 5’-CCAGCTCCAAGAAAGGACGA,R 5’- CGCCCTGTAGGTGAGGTTGAT; *Dhhc3* F 5’- TTACCTAGATGACGTGGGGC,R 5’- AAACGATGACAAAGCCCAGT; *Dhhc7* F 5’-GGTTGGTTCCAGCACACTCT,R 5’- AGAGGAAGAGGATGCCATTG; *Il1a* F 5’- GCACCTTACACCTACCAGAGT,R 5’-AAACTTCTGCCTGACGAGCTT; *Hprt1* F 5′- GATTAGCGATGATGAACCAGGTT, R 5′-CCTCCCATCTCCTTCATCACA; *Ccl2* F 5′- CACTCACCTGCTGCTACTCA, R 5′-GCTTGGTGACAAAAACTACAGC; *CCL2* F 5′-TCTCGCCTCCAGCATGAAAG, R 5′-GGCATTGATTGCATCTGGCT;*Hif1a* F 5′- ACAAGTCACCACAGGACAG, R 5′-AGGGAGAAAATCAAGTCG

### Flow Cytometry

Mice were euthanized, and then buccal mucosal tissue was isolated by first pinning the mouse to a platform and then using scissors to make a midline cut in the bottom lip. Through this cut, scissors were used to separate the skin and underlying muscle around the neck. The mandibular bone was separated by making a cut between the mandibular incisors using scissors. The mandibular incisors were pinned back to expose the entire oral cavity. A razor blade was used to separate the cheek skin from the buccal mucosal tissue. A cut along the maxillary teeth and the mandibular teeth was done to free the buccal mucosal tissue, which was then placed in 3 mL of incomplete RPMI media with L-glutamine. In a tissue culture hood, the tissue was macerated into small pieces using razor blades and then incubated with 5 mL of 0.25% Trypsin in a 37°C incubator for 5 minutes. To quench trypsinization of the tissue, 10 mL of complete RMPI (RPMI media with L-glutamine containing 20% heat-inactivated FBS and 1% penicillin/streptomycin). The RPMI media and the tissue were transferred into a 70 μm strainer and then mashed against the strainer using a syringe plunger. The flow through was collected and then centrifuged, where red blood cells were lysed using 5 mL of ACK lysis buffer for 5 minutes at room temperature. The lysis was stopped by adding 20 mL of sterile PBS buffer, and then the cell suspension was centrifuged. Cells were washed in sterile PBS and then seeded in a 96-U bottom plate at a density of approximately one million cells per well. The plate was centrifuged, and the pelletized cells were resuspended in 100 µL of Zombie Aqua viability dye (423101, BioLegend), at a 1:1000 dilution in sterile PBS, and incubated in the dark at room temperature for 30 minutes. Cells were washed in PBS once and then again in ice-cold FACS buffer (PBS with 2% FBS, 0.1% NaN_3_, and 1 mM of EDTA). To prevent non-specific binding, the cells were incubated with an Fc blocker (anti-CD16/32) (clone 93, BioLegend) for 10 minutes on ice and in the dark. The cells were then stained using the following antibodies: anti-TCRβ (clone H57-597, BioLegend), anti-CD4 (clone RM4-5, BioLegend), anti-PD-1 (clone 29F1A12, BioLegend), anti-Tim-3 (clone RMT3-23, BioLegend), anti-TCRγ/δ (clone GL3, BioLegend), anti-CD3 (clone 17A2, BioLegend), anti-CD11b (clone M1/70, BioLegend), anti-CD206 (clone C068-C2, BioLegend), anti-CD45 (clone 30-F11, BioLegend), anti-CD8 (clone 53-6.7, BioLegend), anti-CD11c (clone N418, BioLegend), anti-Gr1 (Ly6g/c) (clone RB6-8C5, BioLegend), anti-Glut1 (clone EPR3915, Abcam). Surface staining was performed for 30 minutes while maintaining the samples on ice and protected from light. Cells were incubated with continuous agitation on a shaker and intermittently mixed via pipetting during all incubation periods to ensure uniform staining.

### Single-cell Immune Profiling

Tamoxifen was topically applied onto the oral mucosa of *K5-CreER;Sox2^+/+^* and *K5-CreER* mice as previously described. Buccal mucosal tissue from all groups was harvested at the same time, and a cell suspension was extracted as previously described. Immune cells were purified using FACS sorting for CD45^+^ live cells. Considering the size of the buccal mucosa and the fact that one side of the mucosa was utilized to confirm OED/HNSCC diagnosis via histological evaluation, we pooled samples from 6-8 mice per group to obtain a sufficient number of immune cells for single-cell analysis. The cells were submitted in 1X PBS with 0.08% BSA for 10X Genomics 3’-single-cell processing and RNA-Seq at a depth of at least 50,000 reads per cell.

To alleviate the dropout effect, we performed imputations using the R package SAVER. We then removed low-quality cells using the following metrics: the number of unique features is less than 200 or greater than 15,000 and the reads from mitochondria genes are greater than 25%. After filtering, we obtained a total of 8,603 high-quality cells, which were integrated using Seurat. To group cells into different clusters, we employed the Shared Nearest Neighbor method using the top 30 principal components (PCs) from the integrated data with a resolution value of 1. Clusters were annotated using the R package SingleR and manual review of the expression of immune marker genes and those that drove the clustering of cells. To estimate the cell development trajectory, we implemented the R package phateR with the following parameters: the number of k nearest neighbors was 5, the decay value was 40, and t was 10.

### Bone marrow-derived macrophages (BMDMs) isolation and differentiation

To investigate the impact of IL-1α priming on myeloid cell responsiveness to STING agonists, BMDMs were generated by harvesting bone marrow from the long bones of the hind limbs of male and female C57BL/6J mice aged 6-8 weeks. Briefly, mice were euthanized and both femurs and tibiae were harvested using sterilized scissors and forceps before being placed in ice-cold PBS. In a biosafety cabinet, connective tissue and muscle were removed, and the marrow cavity was exposed by cutting off both ends of the bone. Bone marrow was flushed out using a prefilled syringe containing ice-cold sterile PBS and a 27 G ½ needle (305109, BD). The flow through was filtered through a 70-μm cell strainer to achieve a single-cell suspension. Cells were centrifuged and then 2 mL of ACK lysis buffer was added for two minutes at room temperature to lyse red blood cells. Sterile PBS was added to neutralize the ACK lysis buffer followed by centrifugation. The cell pellet was resuspended in 5 mL of BMDM differentiation media (RMPI-1640 with L-glutamine containing 20% heat inactivated FBS, 30% L-929 cell supernatant, and 1% penicillin/streptomycin). The cell suspension was split into four 100 mm tissue culture-treated petri dishes each containing a final volume of 10 mL. Three days post cell seeding, 10 mL of differentiation media was added.

### Myeloid derived suppressor cells (MDSCs) isolation and differentiation

MDSCs were generated by first harvesting long bones and subsequently collecting bone marrow, as previously described. The downstream processing of the cell suspension followed a previously published protocol. In brief, once a single cell suspension was obtained from the bone marrow, 3 million cells were seeded into a 100 mm petri dish containing a final volume of 10 mL of differentiation media (RPMI-1640 with L-glutamine containing 10% heat-inactivated FBS, 10% L929 cell supernatant, 1% penicillin/streptomycin, and 50 µM of β-mercaptoethanol). The cells were cultured at 37°C in a humidified incubator with 5-7% CO_2_ for three days. On the third day, granulocytic and monocytic cells appeared as the main population by FACS analysis and were considered licensed-MDSCs, which require incubation with 100 U/mL IFN-γ and 0.1 µg/mL LPS for 16 hours to become activated-MDSCs.

### MDSC T-cell Suppression Assay

To assess the impact of IL-1α priming on the suppressive function of MDSCs on CD8^+^ T cell proliferation, MDSCs were cultured, following previously described methods, with CFSE-stained CD8^+^ T cells. The dilution of CFSE was then used to determine the level of proliferation. In brief, spleens were harvested from 6-8 week-old male and female C57BL/6J mice and disrupted by pressing them against a 70 µm cell strainer. The resulting single-cell suspension was centrifuged and resuspended for further processing using the EasySep Mouse CD8+ T Cell Isolation Kit. The enriched cell suspension was then thoroughly resuspended in 1 mL of RPMI- 1640 with L-glutamine and 10% heat-inactivated FBS, achieving a final cell concentration ranging from 0.5 × 10^6 to 10^7 cells/mL. To label the CD8^+^ T cells with CFSE dye, the cell suspension was transferred to a fresh, non-wetted tube. To prevent premature mixing with the CFSE solution, the tube was laid horizontally while the CFSE dye was being prepared. One vial of CFSE dye (Invitrogen, C34554) was resuspended in 18 µL of DMSO. Next, 1.1 µL of the stock solution was added to a 110 µL droplet of PBS, which was then placed on the side of the tube. The tube was quickly inverted and vortexed to obtain a uniform solution, and then incubated at room temperature for five minutes while being protected from light. The cells were washed in PBS containing 5% heat-inactivated FBS.

### Acyl-PEG Exchange (APE) Assay

To measure protein palmitoylation, an APE assay protocol was adapted from a previous study^32^. THP-1 monocytes were primed with 10 ng/mL of recombinant mouse IL-1α (I5396, Sigma-Aldrich) for 16 hours. The cells were washed in sterile PBS and subsequently cultured in a complete RMPI medium for four days without any additional IL-1α exposure. Following this incubation period, the cells were stimulated with a STING agonist, 10μg/mL of 2’3’-cGAMP (tlrl-nacga23-1, InvivoGen). The 2’3’-cGAMP was introduced into the culture medium for a duration of 30 minutes, along with human albumin as a carrier. The cells were then collected by centrifugation and then washed two times in sterile PBS before adding 150 μL of chilled lysis buffer which included the following: 50 mM triethanolamine, 4% SDS, 1× protease inhibitor cocktail (11836170001, Roche), 1500 U/mL benzonase (E1014, Sigma), and 5mM PMSF (P7626, Sigma). To inhibit the activity of the benzonase, 5 mM EDTA was added, and protein concentration was measured using a BCA protein assay (Cat. 23225, Thermo Scientific).

Disulfide bonds were cleaved by treating samples with 200 mM TCEP for 30 minutes at room temperature. Then, exposed cysteine residues were capped by incubating the samples with 25 mM of NEM for 2 hours at room temperature. A methanol-chloroform-distilled water precipitation was repeated three times to isolate a protein pellet. The protein pellet was resuspended in TEA buffer supplemented with 4% SDS and 4 mM EDTA, then split into two aliquots to establish hydroxylamine (NH_2_OH) treatment conditions. One aliquot was treated with NH_2_OH and incubated for one hour at room temperature with constant rotation to facilitate the cleavage of thioester bonds between cysteine residues and fatty acids. The second aliquot, which did not receive NH2OH treatment, acted as the negative control and was incubated under identical conditions. To remove NH_2_OH, a methanol-chloroform precipitation step was performed on both the treated and control samples. The protein pellet was resuspended in TEA buffer containing 4% SDS and 4mM EDTA. The addition of a 5 kDa mass tag to the previously coupled cysteine residues was accomplished by incubating the protein solution with 1 mM mPEG-Mal for two hours at room temperature and under constant rotation. A final methanol-chloroform-distilled water precipitation was performed. Samples were resuspended in 1× Laemmli buffer, boiled at 95°C for 5 minutes, and then separated by SDS/PAGE and analyzed by immunoblotting. Lysates were loaded into a 4-20% tris-glycine, 1.0 mm polyacrylamide gel (XP04200BOX, Invitrogen). Proteins from lysates were blotted using the following antibody: Sting antibody (13647S, RRID: AB_2732796, Cell Signaling Technology).

### Quantification and statistical analysis

To compare statistical differences between two independent groups, a two-tailed Student’s t-test was used. For comparisons made between more than two groups, a two-way ANOVA followed by Šidák’s multiple comparisons test was utilized. For tumor volume comparisons, a generalized estimating equation was performed. For all Figures, a statistical significance was indicated according to the following scale:*p < 0.05; **p< 0.01; ***p< 0.001; and ****p< 0.0001. Data in the graphs are expressed as mean ± SEM.

### Study Approval

All animal procedures were performed in accordance with the protocol approved by the IACUC at the M.D. Anderson Cancer Center (00002478-RN00) and the University of Michigan (PRO00010232). The use of human specimens was approved by the M.D. Anderson Cancer Center Institutional Review Board (2024-0364) and the University of Michigan IRB (HUM00042189 and HUM00113038).

## Acknowledgments

This work is supported by NIH grants R01 DE026728, R01 DE030691, R01 DE031951, U01 DE029255, U01 DE033330, T32 DE007057 and F31DE033922.

## Author Contributions

Author Contributions: H.F.T., W.G., Z.R.F., Z.L., Y.H., W.C., performed experiments, data processing, and figure generation. J.Q. provided key animal resources. K.O., J.K., S.R. and A.M. preformed cell lines generation and histology. S.F., M.L., F.N. and E.L. provided clinical resources and advice. Y.W., Y.X. and Y.L.L. analyzed bulk RNA-seq and single cell RNA-seq. H.F.T., J.J.M. and Y.L.L. designed the project with significant intellectual input from S.D., P.J.P., J.E.N. T.D.W., and H.W.. H.F.T., W.G. and Y.L.L. wrote the manuscript and all authors edited the manuscript and have read and agreed to its contents.

## Figure Legends

**Supplemental Figure 1.**
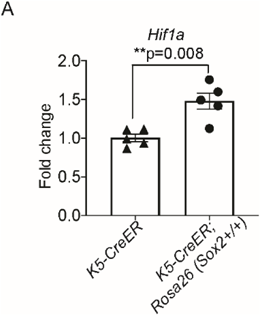
Glycolysis signature is increased in Sox2-driven cancer. (A) Increased *Hif1a* expression was observed in the buccal mucosa of tamoxifen-treated *K5-CreER;Sox2^+/+^* mice (n=5) compared to control *K5-CreER* mice (n=5). Mice were euthanized about 3 weeks post tamoxifen treatment, and total RNA was isolated from buccal mucosa treated with 40 mg/mL tamoxifen for five consecutive days. Expression of *Hif1a* was assessed via qPCR. Values represent mean ± SEM of three biological replicates per sample. Comparisons between the two groups were made using a two-tailed unpaired t-test (**p<0.01).

**Supplemental Figure 2.**
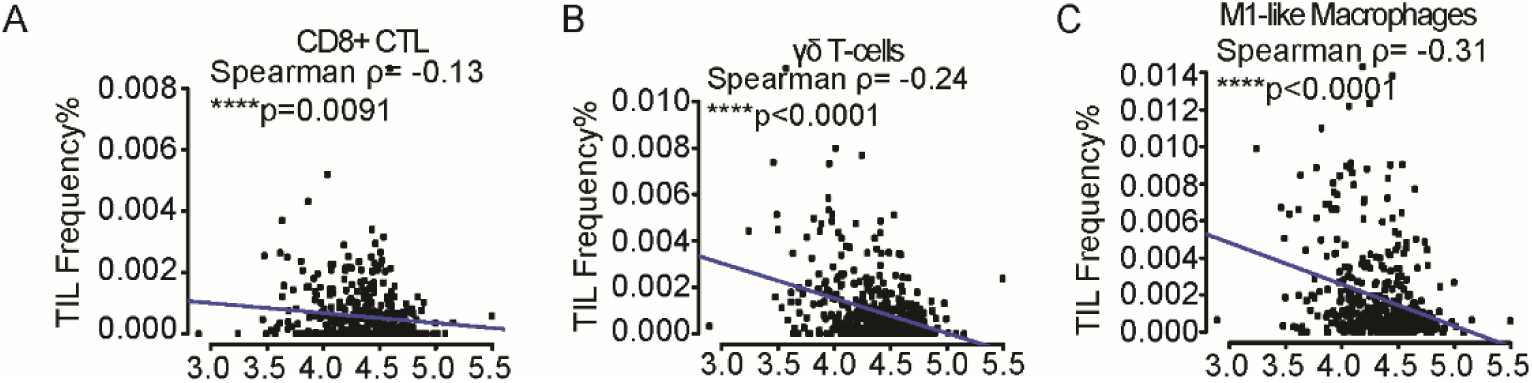
Correlation of *SLC2A1* Expression with Immune Subsets in HPV^-^ HNSCC. *SLC2A1* expression levels in human Head and Neck Squamous Cell Carcinoma (HNSCC) from TCGA were correlated with the abundance of CD8^+^ CTL (A), γδ T-cells (B), and M1-like macrophages (C) in HPV^-^ tumors.

**Supplemental Figure 3.**
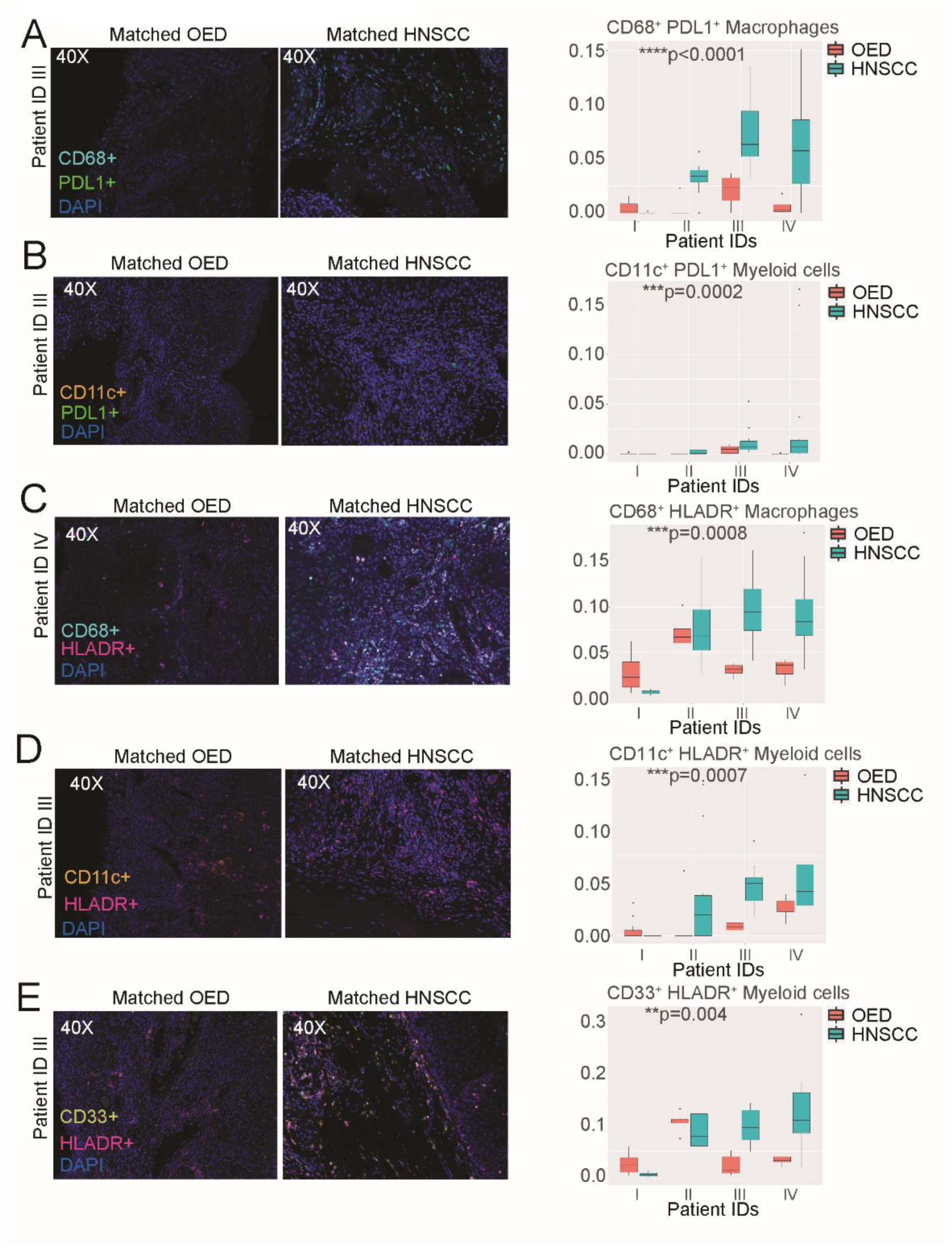
The immune landscape shifts as OED transforms to HNSCC. (A-E) Representative multispectral immunofluorescence images (MSI) of tumor-infiltrating myeloid and lymphoid cells from paired OED and HNSCC tissue specimens. Images were captured using the Akoya Biosciences Vectra Polaris scanner, and regions of interest were selected with Akoya Phenochart software. Data analysis was performed using the Akoya phenoptrReports package in RStudio, followed by a two-sample Wilcoxon test comparing mean expression levels between matched OED and HNSCC stages for each patient. Representative MSI images of CD68^+^ PDL1^+^ macrophages (A), CD11c^+^ PDL1^+^ myeloid cells (B), CD68^+^ HLADR^+^ macrophages (C), CD11c^+^ HLADR^+^ myeloid cells (D), CD33 ^+^ HLADR ^+^ myeloid cells (E) in matched OED/HNSCC tissues are shown. Results were considered significant if the p-value was less than 0.05.

**Supplemental Figure 4.**
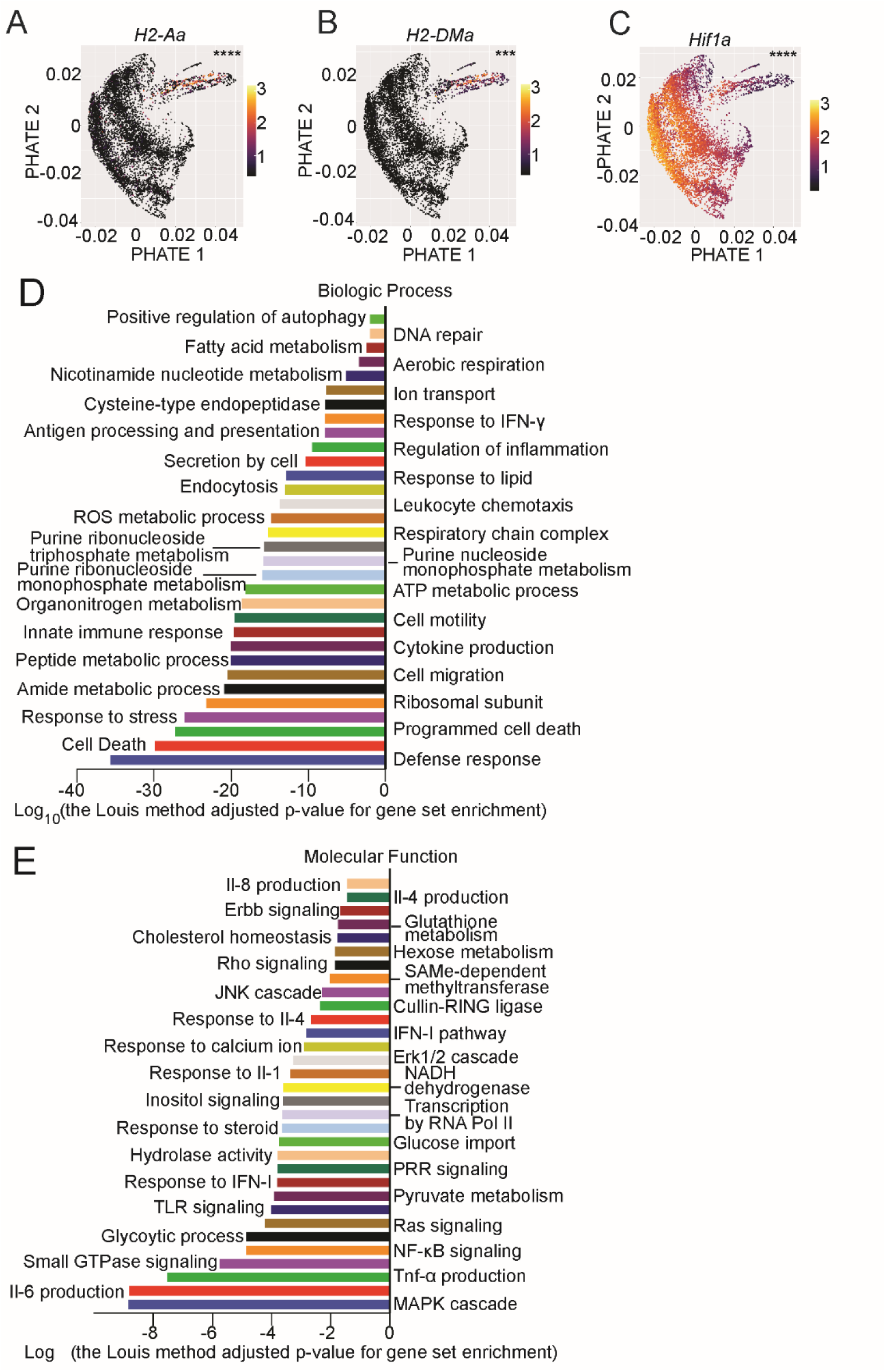
The evolution of myeloid cells during Sox2-driven HNSCC initiation. (A-C) Buccal mucosal tissue from *K5-CreER^+^;Sox2^+/+^* mice at 2-weeks and 4-weeks post tamoxifen administration, and K5-CreER^+^ control mice were harvested, CD45^+^ immune cells were isolated by FACS, and subjected to scRNA-Seq. PHATE trajectory analysis of myeloid cells was performed. The expression of *H2-Aa* (A), *H2-DM*a (B), and *Hif1a* (C) are highlighted. (D-E) GSEA comparing normal mucosal-resident and intra-lesional myeloid cells was performed using the biologic process and molecular function gene sets.

**Supplemental Figure 5.**
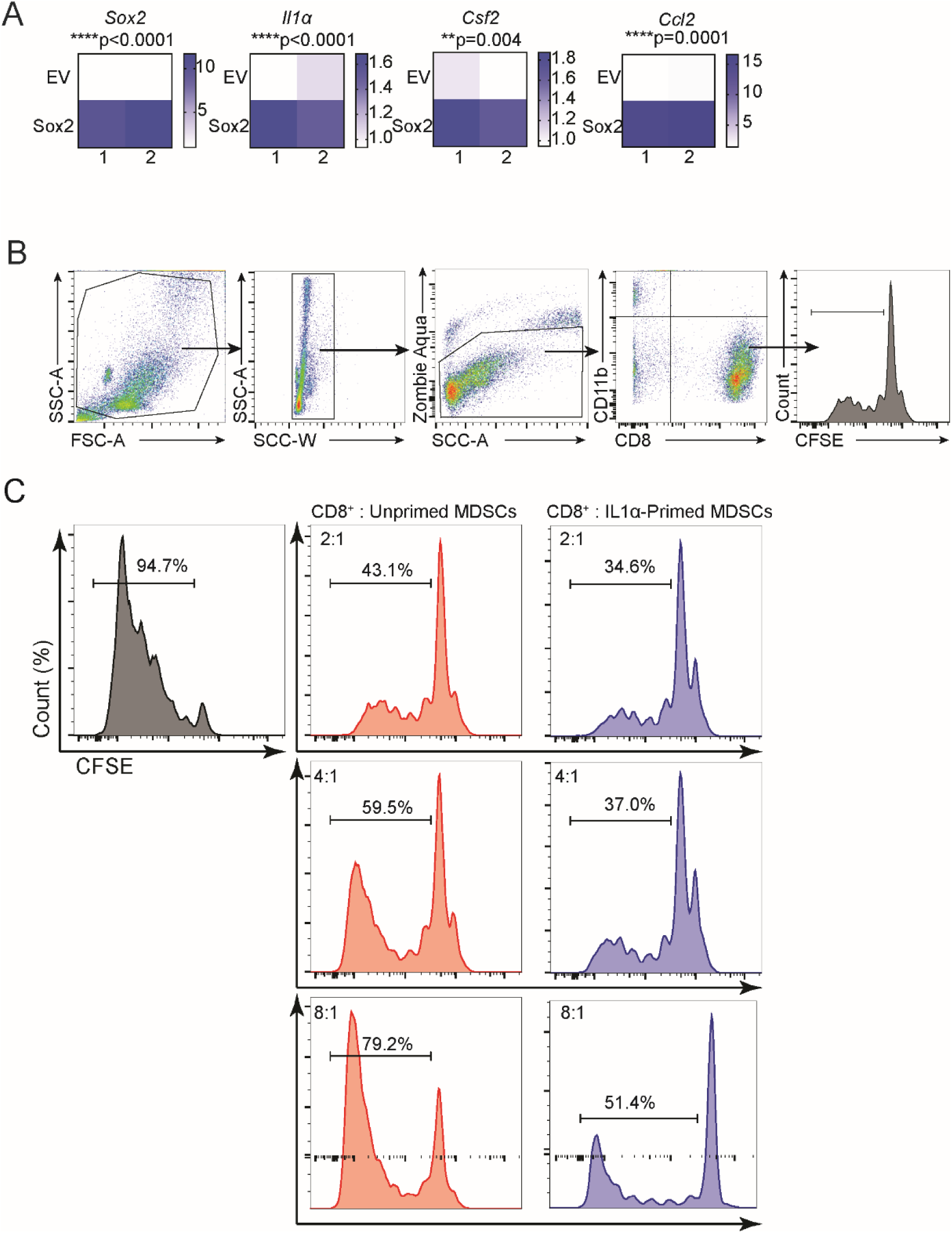
IL1α-primed MDSC are even more suppressive of T-cell activation. (A) Empty vector control and Sox2-expressing MOC2-E6/E7 cells were subjected to bulk RNA-Seq. Heatmaps show the significant difference in the expression levels of *Sox2, Il1a, Csf2,* and *Ccl2* between the two groups. (B-C) MDSCs were primed with 10 ng/mL IL-1α or PBS control for 16 hours, rested, and then activated with IFN-γ and LPS. CD8^+^ T-cells, isolated from spleens and labeled with CFSE, were co-cultured with the MDSCs, IL-2, and beads for 4 days. Cells were gated for single, live, CD45^+^, and further gated on CD11b^-^ CD8^+^ T cells. Representative scatterplots show CFSE dilution in CD8^+^ T cells. A two-tailed unpaired t-test was used for comparisons (*p<0.05).

**Supplemental Table S1**

The table shows the most differentially expressed genes between the myeloid cells from oral normal mucosa and those from transformed epithelial tissues at the two-week and four-week time points.

**Supplemental Table S2**

The table shows the top 200 most differentially expressed genes between EV-MOC2-E6/E7 and SOX2-expressing MOC2-E6/E7 cells.

